# Temporal confounds emulate multivariate fMRI measures of perceptual learning

**DOI:** 10.1101/2025.08.12.669686

**Authors:** Georgia Milne, Kim Staeubli, Hugo Chow-Wing-Bom, John Greenwood, Peter Kok, Roni Maimon Mor, Tessa Dekker

## Abstract

Human perception inherently involves learning. We can experience the same stimulus completely differently depending on our prior knowledge. Understanding the neural basis of perception therefore requires measurements that capture these temporal dynamics. Multi-voxel pattern analysis (MVPA) approaches are widely used to characterise changes in neural representations over time. A popular example is the perceptual reorganisation paradigm, which investigates the neural correlates of enhanced recognition of distorted images after cueing with the undistorted version. Studies typically report increased representational similarity of these two images in early visual cortex. However, as these paradigms include an inherent ordering of stimuli that precludes trial randomisation or counterbalancing, they are vulnerable to temporal confounds common to fMRI. Here, we investigate how these confounds could influence current understanding of perceptual reorganisation. We tested different perceptual reorganisation paradigm designs derived from published fMRI studies, and found substantial design-driven order effects at the single-subject level for all paradigms. For certain designs, these effects artificially amplified neural indices of perceptual reorganisation at both the single-subject and group levels, emulating widespread signatures of perceptual reorganisation across the brain. To disentangle perceptual learning processes from measurement artefacts, we recommend (i) selecting designs that minimise the effect of stimulus order confounds on contrasts of interest, (ii) correcting for these confounds, and (iii) confirming results are perceptually driven via negative controls, e.g., stimuli or brain areas not expected to produce perceptual effects. Our work demonstrates how current understanding of perceptual learning mechanisms based on multivariate neuroimaging approaches could be influenced by non-obvious design confounds that misdirect interpretations towards distributed neural processing, and offers practical solutions to address this.

## Introduction

Sensory representations are constantly updated as we learn. For example, long-term exposure to a particular species of bird can make its details more salient, or hearing a friend’s voice can cause you to instantly recognise someone you previously thought was a stranger. In cognitive neuroscience, these perceptual dynamics are often studied with cueing tasks (Carrasco, 2018; Schacter & Buckner, 1998), where cues that manipulate knowledge or attention alter neural responses to subsequent stimuli. With the emergence of multivariate fMRI techniques, this is typically measured via representational similarity analysis (RSA), where variations in neural representations to different stimuli are assessed by correlating their multi-voxel responses. While these measures are informative and robust to various univariate signal fluctuation confounds (Kriegeskorte, 2008; Roth & Merriam, 2023), structured noise in the temporal domain can affect their results (Alink et al., 2015; Cai et al., 2019; Mumford et al., 2014). This could be particularly problematic for learning paradigms, as they often require stimuli to be presented in a specific sequence (e.g., pre-cueing target, cue, post-cueing target), making full randomisation or counterbalancing experimental order impossible.

A popular cueing task uses two-tone images to study perceptual reorganisation (Chang et al., 2016; Dolan et al., 1997; Flounders et al., 2019; González-García et al., 2018; González-García & He, 2021; Hsieh et al., 2010; Milne et al., 2024; van Loon et al., 2016; Yoon et al., 2007). In this phenomenon, an unrecognisable two-tone image, created by smoothing and contrast-binarising a greyscale photo (see Figure 1A), becomes recognisable only after exposure to the original photo version (Mooney, 1957). Perceptual reorganisation can therefore be robustly triggered by the sequential presentation of a two-tone image, its corresponding photo cue, and the same two-tone again (Ramachandran, V S, 1994). Because newly acquired knowledge can provoke distinct perceptual experiences in response to two identical images, this offers a well-controlled paradigm for measuring top-down modulation of sensory inputs across the brain.

**Figure 1:**
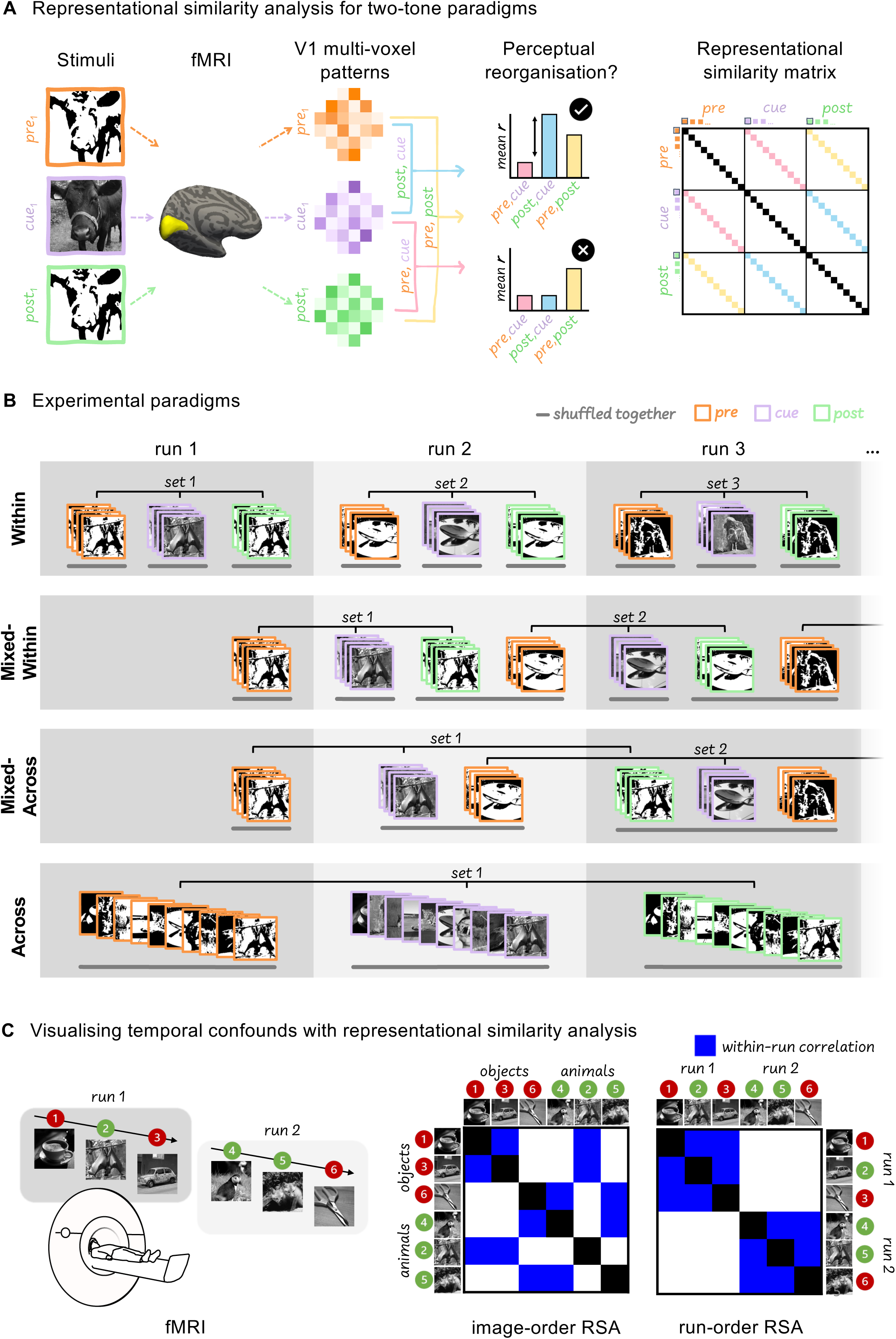
Perceptual reorganisation paradigms and analyses. **A** Experimental procedure and Representation Similarity Matrix (RSM) construction for measuring perceptual reorganisation with fMRI. **B** Paradigm sequences of the four experimental designs tested shown for the first three fMRI runs of each design (total number of runs: Within, five runs; Mixed-Within, six runs; Mixed-Across, seven runs; Across, six runs. Each image is shown in ‘Pre’ condition (orange borders), ‘Cue’ condition (lilac borders) and ‘Post’ condition (green borders). Groups of images shuffled together are underlined with grey bars. **C** Temporally driven pattern correlations, e.g. positive correlations of trials within the same run (within-run correlations), are obscured by conventional RSA methods that order data according to image identity (image-order RSA, left matrix), but can be visualised by ordering the same data by the sequence in which stimuli were presented across the experiment (run-order RSA, right matrix).

RSA can reveal whether two-tone images are represented more similarly to their photo versions after cueing, in line with the phenomenological experience of perceptual reorganisation (Figure 1A). Hsieh and colleagues were the first to identify this neural marker of perceptual reorganisation (Hsieh et al., 2010). They observed perceptual reorganisation in the Lateral Occipital Complex (LOC), a higher-order area known to play a critical role in object recognition (Malach et al., 1995), but also in early visual cortex, suggesting widespread knowledge-based modulation of neural representations (Hsieh et al., 2010). Van Loon and colleagues later extended these findings, observing widespread effects of perceptual reorganisation from primary visual cortex (V1) to posterior fusiform LOC, which were diminished in lower-order areas when NMDA-mediated feedback processing was inhibited (van Loon et al., 2016). More recently, Gonzáles-García and colleagues replicated findings of distributed knowledge-driven processing, and further identified classically ‘non-visual’ areas, such as the frontoparietal (FPN) and default-mode (DMN) networks, as encoders of perceptual reorganisation (González-García et al., 2018; González-García & He, 2021). Strikingly, these studies find that classically ‘input-driven’ areas, such as V1, represent cued two-tones more similarly to their photo versions than to the same two-tone viewed before cueing (González-García et al., 2018; Hsieh et al., 2010; van Loon et al., 2016). While top-down processing has previously been found to influence representations throughout the visual hierarchy in other perceptual tasks (Kok et al., 2012; Lamme & Roelfsema, 2000; Poltoratski & Tong, 2020; Self et al., 2013; Yon et al., 2022), the findings from perceptual reorganisation studies extend this considerably by reporting effects of prior knowledge that are widespread across the cortex, and that can dominate incoming visual input.

The above findings offer compelling insights into long-standing neuroscientific debates about the distribution of conscious perceptual processing and attention, leveraging a range of paradigm designs to reach corroborating conclusions about the strength and distribution of top-down neural signals. However, the designs used share a feature that is common to all perceptual reorganisation paradigms: uncued two-tones (‘Pre’ condition) must always precede their corresponding photo versions (‘Cue’ condition), which in turn must always precede the corresponding cued two-tones (‘Post’ condition, Figure 1A). As this presentation order is necessary for triggering and measuring perceptual reorganisation, it results in a temporal consistency across subjects that cannot be avoided by trial randomisation and counterbalancing. This raises questions about the potential influence of temporal artefacts on RSA measures of perceptual reorganisation, which may not be captured using conventional methods of reporting these data (Figure 1C).

In fMRI, structured noise can arise from both physiological (e.g. subject motion, breathing, heart-rate and attentional drift) and methodological (e.g. thermal fluctuations, scanner instability, equipment radio frequency signals, and preprocessing and analysis artefacts) sources, and act at multiple temporal levels, i.e., between scanning sessions, within a single session, between fMRI runs, and within single runs (Alink et al., 2015; Cai et al., 2019; Mumford et al., 2014; Zarahn et al., 1997). The combined impact of structured noise on multi-voxel patterns may therefore be complex, and variable across subjects, set-ups and paradigm designs. To investigate this issue, we tested four designs of two-tone paradigms based on examples in the literature, and characterised the impacts of temporal effects at the level of individual participants in each design.

The paradigms differed in how stimulus conditions (Pre, Cue and Post) were distributed across the experiment, and should therefore vary in their sensitivity to structured noise across different temporal levels. In an across-run design (Hsieh et al., 2010; van Loon et al., 2016), separate fMRI runs are dedicated to each stimulus condition, allowing trials to be fully randomised within each run without disrupting the required order of conditions (‘Across’ design; Figure 1B, bottom row). In a ‘Mixed-Within’ design (Flounders et al., 2019; González-García et al., 2018; González-García & He, 2021), Pre images are presented in a separate run to the corresponding Cue and Post images, which are shown in the following run alongside new Pre images. This design counterbalances stimulus conditions more evenly across the experiment, but unsymmetrically across fMRI runs (Figure 1B, second row). We also included two further paradigm designs that offer logical alternatives for managing temporal order constraints. In a Mixed-Across design, a single run can contain all three stimulus conditions, but Pre, Cue and Post conditions of the same images are always separated across three consecutive runs, allowing for randomisation of all images presented within a run (Figure 1B, third row). Finally, a Within-run design presents all three conditions of a set of images within a single run, meaning trials can only be randomised within subsections of each run, but effect sizes may be enhanced by avoiding across-run signal variability (Figure 1B, first row).

For each of the above four designs, we presented the same 20 images of natural and man-made scenes, and measured perceptual reorganisation behaviourally and cortically at the single-subject level. Our cortical regions of interest included primary visual cortex (V1) and primary somatosensory cortex (S1) as a control region. While previous studies have reported strong patterns of perceptual reorganisation in V1 (González-García et al., 2018; González-García & He, 2021; van Loon et al., 2016), S1, as a typically non-visual region, should not encode perceptual reorganisation of visual stimuli, but may show design-driven artefacts. Using conventional RSA measures, we find that some paradigms result in strong neural patterns of perceptual reorganisation across both V1 and S1, while others do not. This leads us to identify a substantial temporal artefact present in all paradigms tested, that confounds conventional measures of perceptual reorganisation depending on the distribution of stimulus conditions across fMRI runs. We further show how even in confounded paradigms, appropriate control analyses can mitigate these artefactually enhanced effects. Critically, without these corrections, the true extent of temporal confounds can be masked by conventional RSA practices (Figure 1C), yet can emulate strong neural patterns of perceptual reorganisation across the cortex. Based on this example from perceptual reorganisation, we recommend a systematic three-step approach to distinguish genuine learning effects from temporal artefacts in multivariate fMRI studies.

## Results

We assessed four fMRI paradigm designs of a two-tone cueing task that included 20 images presented in three conditions: as a naively viewed two-tone (Pre), followed by the original photo (Cue), and the same two-tone viewed after cueing (Post). The paradigms varied in how these conditions were organised across fMRI runs (Figure 1B); where the three conditions of any given image were presented in either a single run (‘Within’ paradigm), two runs (‘Mixed-Within’ paradigm), or three runs (‘Mixed-Across’ paradigm and ‘Across’ paradigms). For all paradigms, image perception was measured via a verbal recognition task completed between fMRI runs, and an animate/inanimate categorisation task completed during each trial. Photo cueing improved two-tone recognition, increasing the average number of correctly identified images from 39% to 90% across all designs (Supplementary 1).

We tested for neural markers of perceptual reorganisation using representational similarity analysis (RSA; Kriegeskorte, 2008). After running a GLM, we used the univariate parameter estimates to calculate the average multi-voxel response pattern elicited by each of the 20 images (Im_1_ to Im_20_) for each of the three stimulus conditions (Pre, Cue, and Post). We restricted these patterns to our two cortical regions of interest (ROIs): primary visual cortex (V1) where substantial perceptual reorganisation has been previously reported, and a control region, primary sensory cortex (S1). Next, we applied multivariate noise normalisation (Walther et al., 2016) and compared all 60 response patterns to each other via pairwise correlations (*pearson’s **r***) to produce image-order representational similarity matrices (RSMs), in which each square represents the similarity of two individual response patterns (group-averaged RSMs shown in Figure 2A, representative single-subject RSMs shown in Supplementary 2). We then calculated the mean similarity for each across-condition image pair in single-subject RSMs (within-image ***r***; Figure 2A).

**Figure 2:**
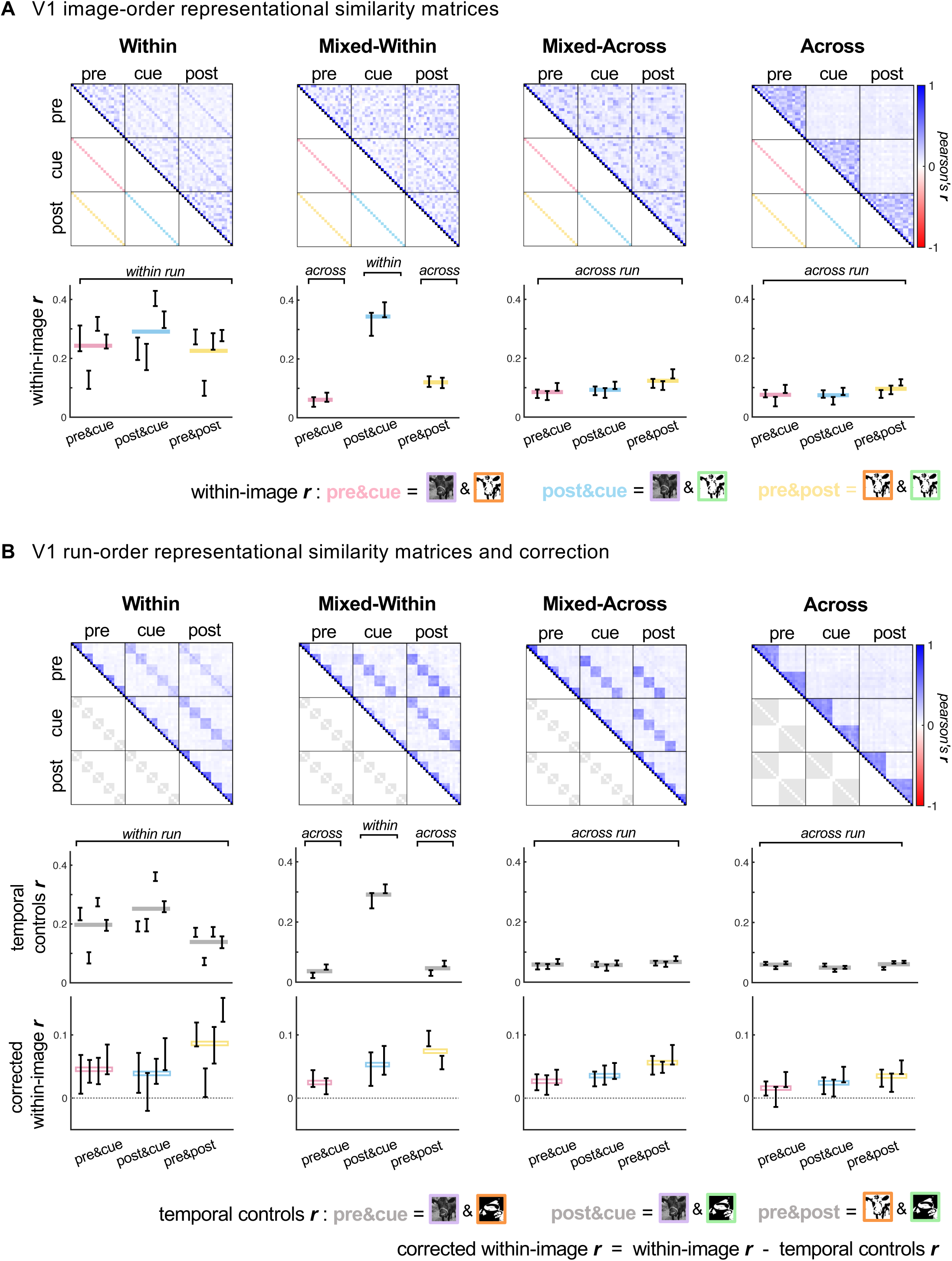
Temporal order effects correlate with RSA-derived measures of perceptual reorganisation in primary visual cortex. **A** Top row: Group-averaged image-order RSMs for primary visual cortex (V1), ordered by stimulus condition (Pre, Cue and Post) and image identity (Im_1_ to Im_20_) for each paradigm group (Within: n = 4, Mixed-Within: n = 2, Mixed-Across: n = 3, Across: n = 3). Correlations of the same image presented in different conditions are represented along RSM off-centre diagonals (pink diagonal for Pre&Cue, blue diagonal for Post&Cue, and yellow diagonal for Pre&Post), where greater representational similarity is expected, denoted by darker blue colours, compared to for different images. Bottom row: Mean within-image similarities for across-condition comparisons in V1. Error bars show bootstrapped 95% confidence intervals for individual subjects, horizontal coloured bars show group averages for each across-condition comparison. **B** Top row: Group-averaged run-order RSMs for V1, ordered by stimulus condition (Pre, Cue and Post) and experiment presentation order. Grey squares indicate temporal controls for each paradigm: pairs of non-corresponding images that appear in the same fMRI runs as the corresponding images in each across-condition comparison. Middle row: Mean similarities of temporal controls for across-condition comparisons in V1. Error bars show bootstrapped 95% confidence intervals for individual subjects, horizontal grey bars show group averages for each across-condition comparison. Bottom row: Correction of mean similarities for across-condition comparisons of corresponding images in V1. For each across-condition comparison, within-image similarities are corrected by subtracting mean similarities of temporal controls within each image set and ROI at the single-subject level. Error bars show bootstrapped 95% confidence intervals for individual subjects, horizontal hollow bars show group averages for each across-condition comparison.

Under perceptual reorganisation, the neural representation of a two-tone image after cueing (Post) should more closely match the representation of its original photo (Cue). Hence, the correlation between cued two-tones and their corresponding photo cues (Post&Cue, blue diagonal) should be higher than for uncued two-tones and their corresponding photo cues (Pre&Cue, pink diagonal). Perceptual reorganisation is therefore defined as Post&Cue > Pre&Cue.

There was a large perceptual reorganisation effect (Post&Cue > Pre&Cue) at the single subject level in the Mixed-Within paradigm (t(19) > 12, Cohen’s *dz* > 2.6, *p* < 10^-9^), a more moderate effect in the Within paradigm (t(19) > 2.4, Cohen’s *dz* > 0.56, *p* < 0.05 for three of four subjects) and a nonsignificant effect in across-run designs (*p* > 0.05 for all subjects; see supplementary 4 for all statistical results). These differences occurred despite all paradigms producing a large behavioural effect (Supplementary 1). Multivariate fMRI measures are highly susceptible to temporal differences in stimulus presentation. For example, combining across all paradigms, correlations of corresponding images across conditions presented within the same run (within-run) were higher than those presented in different runs (between-run; Figure 2A; linear mixed model: β(within/between) = 0.276, SE = 0.013, t(714) = 21.79, p < 0.001, 95% CI [0.251, 0.301]). This is especially problematic in the mixed-within paradigm, where the Post&Cue comparison is made within a run, and the Pre&Cue comparison is made across runs, as this confounds the key contrast for perceptual reorganisation with temporal order structure in the BOLD signal. However, temporal artefacts can arise at multiple timescales across an experiment, for example in addition to within-versus-across-run differences, correlations may drift within a run or across the session.

Therefore, all four paradigms can be affected by temporal consistencies in stimulus presentation sequences. We therefore next explored whether the observed perceptual reorganisation effects also exist for pairwise image comparisons with equivalent temporal relationships, but that should not generate a cognitive effect of perceptual reorganisation: firstly, in a control cortical ROI (primary somatosensory cortex [S1]) and secondly, in non-corresponding images.

Because perceptual reorganisation reflects knowledge-based modulation of visual representations, we would not expect to find neural markers of this effect in primary somatosensory cortex (S1). However, in this non-visual ROI, we nevertheless found strong patterns of perceptual reorganisation in the Within and Mixed-Within paradigms (Figure 3A). Mirroring findings from V1, Post&Cue correlations exceeded Pre&Cue correlations in Mixed-Within (t(19) > 11, Cohen’s *dz >* 2.3, *p* < 10^-8^) and Within designs (t(19) > 4.2, Cohen’s *dz* > 0.95, *p* < 0.001 for three of four subjects; see Supplementary 4 for all statistical results). The presence of perceptual reorganisation markers in multivariate fMRI in non-visual regions, where they would not be expected, suggests a susceptibility of this approach to methodological artefacts that could falsely emulate neurocognitive learning effects.

**Figure 3:**
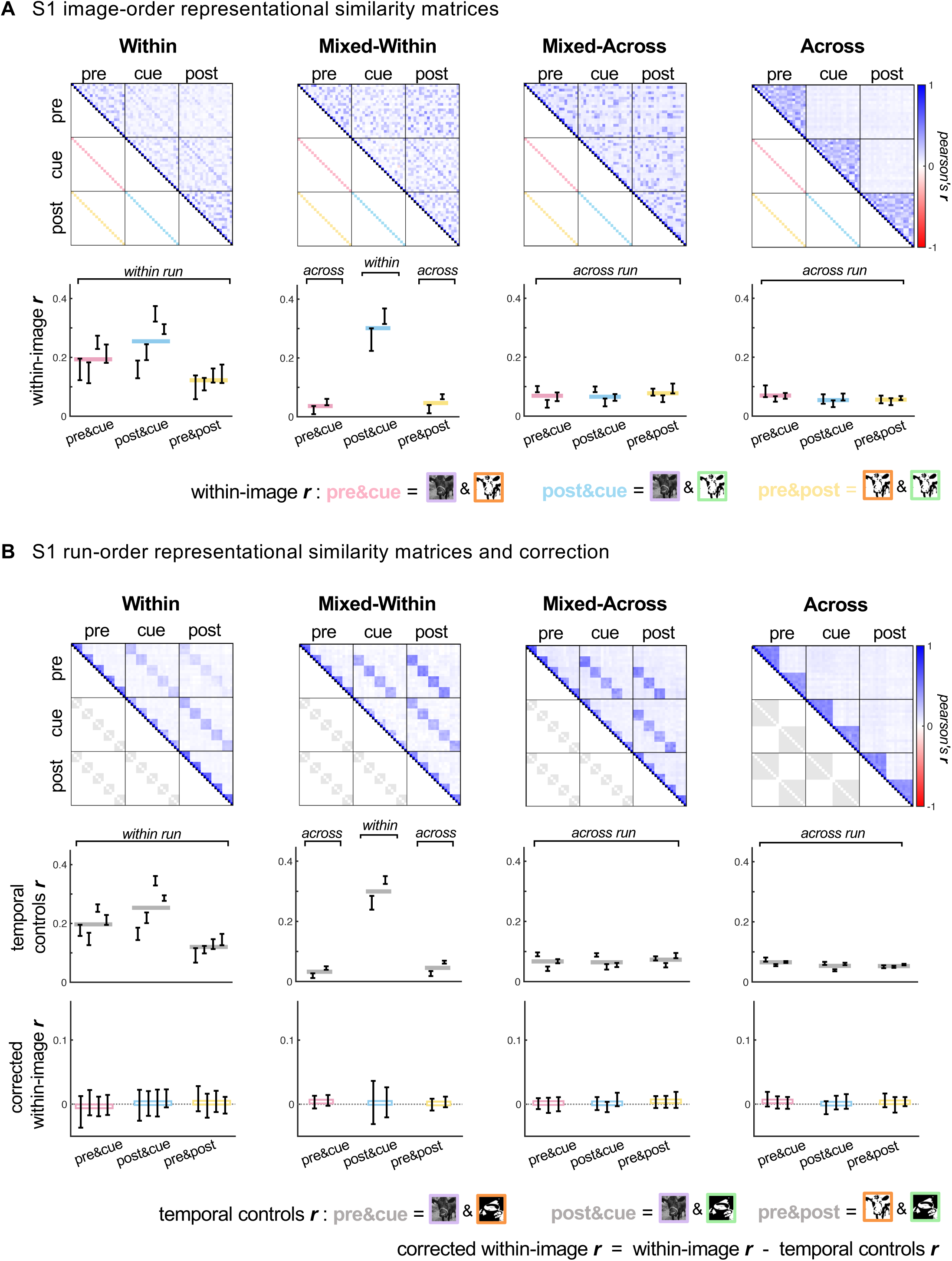
Temporal order effects extend beyond visual cortex, emulating perceptual reorganisation in S1. **A** Top row: Group-averaged image-order RSMs for primary somatosensory cortex (S1), ordered by stimulus condition (Pre, Cue and Post) and image identity (Im_1_ to Im_20_) for each paradigm group (Within: n = 4, Mixed-Within: n = 2, Mixed-Across: n = 3, Across: n = 3). Correlations of the same image presented in different conditions are represented along RSM off-centre diagonals (pink diagonal for Pre&Cue, blue diagonal for Post&Cue, and yellow diagonal for Pre&Post), where greater representational similarity is expected, denoted by darker blue colours, compared to for different images. Bottom row: Mean within-image similarities for across-condition comparisons in S1. Error bars show bootstrapped 95% confidence intervals for individual subjects, horizontal coloured bars show group averages for each across-condition comparison. **B** Top row: Group-averaged run-order RSMs for S1, ordered by stimulus condition (Pre, Cue and Post) and experiment presentation order. Grey squares indicate temporal controls for each paradigm: pairs of non-corresponding images that appear in the same fMRI runs as the corresponding images in each across-condition comparison. Middle row: Mean similarities of temporal controls for across-condition comparisons in S1. Error bars show bootstrapped 95% confidence intervals for individual subjects, horizontal grey bars show group averages for each across-condition comparison. Bottom row: Correction of mean similarities for across-condition comparisons of corresponding images in S1. For each across-condition comparison, within-image similarities are corrected by subtracting mean similarities of temporal controls within each image set and ROI at the single-subject level. Error bars show bootstrapped 95% confidence intervals for individual subjects, horizontal hollow bars show group averages for each across-condition comparison.

We therefore considered how temporal order structure in the BOLD signal may impact representational similarity measures of perceptual reorganisation across the different paradigms we tested. To explore this, we constructed additional ‘run-order RSMs’ (see Methods; Figure 2B) that are ordered according to when each image was presented across the experiment - which differed for every subject due to trial pseudorandomisation. Rows and columns of RSMs are arranged by stimulus condition (Pre, Cue and Post), and within each condition are traditionally arranged in a constant pre-defined order of images across all subjects, regardless of the order in which the images were presented. In a run-order RSM, images are arranged by the unique order in which they were presented to each subject. So the first rows will include the first shown images, but the identity of these images would be different for each subject. Notably, run-order RSMs still display correlations of corresponding images along the off-centre diagonals, but in addition will visualise any temporally-driven correlation structures that would otherwise be obscured by image-order RSMs (see Figure 1C).

In addition to the increased correlations between corresponding images across different conditions previously visible in image-order RSMs, the run-order RSMs also reveal temporal correlation structure unrelated to stimulus identity. One striking feature is that multi-voxel patterns from any two images presented in the same fMRI run are more correlated than those presented in different runs in all paradigms - resulting in dark blue blocks for any comparisons made within a run. This is a well-known feature of fMRI BOLD timeseries measures (Mumford et al., 2014), more visible after correction for common noise across the session but also clearly present in our unnormalised data (Supplementary 3). This contextualises the earlier reported finding of increased correlations for corresponding images within-versus across-run. Run-order RSMs are useful for visualising how the arrangement of conditions across fMRI runs can confound perceptual learning indices (here Pre&Cue>Post&Cue) with substantial temporal correlation structure in the timeseries. For example, it becomes clear how within-run vs across-run correlation structure would artifactually inflate the perceptual reorganisation effect in our Mixed-Within design: Post&Cue correlations of corresponding images, which always fall within a run and so are artefactually enhanced, are compared with Pre&Cue correlations that always fall across different runs.

To assess the impact of temporal artefacts on RSA measures of perceptual reorganisation, we used a control measure that isolates the effects of temporal experimental structure on multi-voxel pattern correlations from those relating to image identity. Specifically, for each comparison of corresponding images across conditions (i.e., Pre&Cue, Post&Cue, and Pre&Post), we calculated the average representational similarity of non-corresponding images with equivalent temporal relationships, and used these as temporal controls (light grey squares in RSMs in Figures 2B and 3B). Because these temporal controls involve images from the same conditions and runs as the main within-image comparisons, they should capture the same temporal correlation structure but not the perceptual learning effects specific to the shared identity of corresponding images. This analysis revealed that in the Within and Mixed-Within designs, temporal-order artefacts produce a correlation structure that mimics a pattern of perceptual reorganisation; Post&Cue similarity is higher than Pre&Cue similarity for temporal controls in both V1 and S1 in these designs (Figure 2B; Mixed-Within: t(59) > 17, Cohen’s *dz* > 2, *p* < 10^-25^; Within: t(59) > 4, Cohen’s *dz* > 0.5, *p* < 10^-4^ for three of four subjects; see Supplementary 5 for all statistical results). This control analysis reveals temporal artefacts with a widespread presence across the cortex.

Next, to isolate image-driven neural effects from temporally driven artefacts, we corrected comparisons of interest by subtracting the mean correlation of temporal controls on a run-by-run basis. This results in a similarity measure that appropriately corrects for the temporal confounds that each image comparison is subject to across multiple timescales, isolating image-specific changes in neural representations (see Methods). At the single-subject level, we found that this correction removed the consistent patterns of perceptual reorganisation in V1, previously present in six individuals, and now in just one out of these six (Figure 2B, bottom panel; see Supplementary 6 for all statistical results). We cannot conclude from these results that V1 image encoding does not perceptually reorganise - as effects may be too small to detect at the single-subject level - but it does mean that the large effect sizes previously observed were erroneous.

Correcting for temporal order confounds effectively normalises the data and makes it possible to combine measures across paradigms. Across all 12 participants, in line with the role of V1 in low-level visual feature encoding, V1 representational similarity was now highest for repeated presentations of the two-tone images before and after cueing (Figure 4, left panel; Pre&Post; paired t-test across subjects, Pre&Post vs. Pre&Cue: t(11) = 4.7, Cohen’s *dz* = 1.4, p < 0.001; Pre&Post vs. Post&Cue: t(11) = 4.2, Cohen’s *dz* = 1.2, p < 0.01). Meanwhile, no significant group-level effect of perceptual reorganisation in V1 remained following this correction (Post&Cue vs. Pre&Cue: t(11) = 1.4, Cohen’s *dz* = 0.42, p > 0.05), although it is possible that subtle cueing modulations in V1 may be detected with more powerful designs. In S1, correcting for temporal artefacts resulted in near-zero correlations across all conditions and paradigms, abolishing any previous evidence of visual perceptual reorganisation in this non-visual brain area at both the single-subject level (Figure 3C, bottom panel; *p* > 0.05 for all comparisons) and the group level (Figure 4, right panel; t(11) < 0.9, Cohen’s *dz* < 0.3, p > 0.05 for all comparisons). Together, this shows that multivariate fMRI measures of perceptual learning could be strongly inflated by non-specific temporal order effects in the data, if not accounted for with appropriate controls.

**Figure 4:**
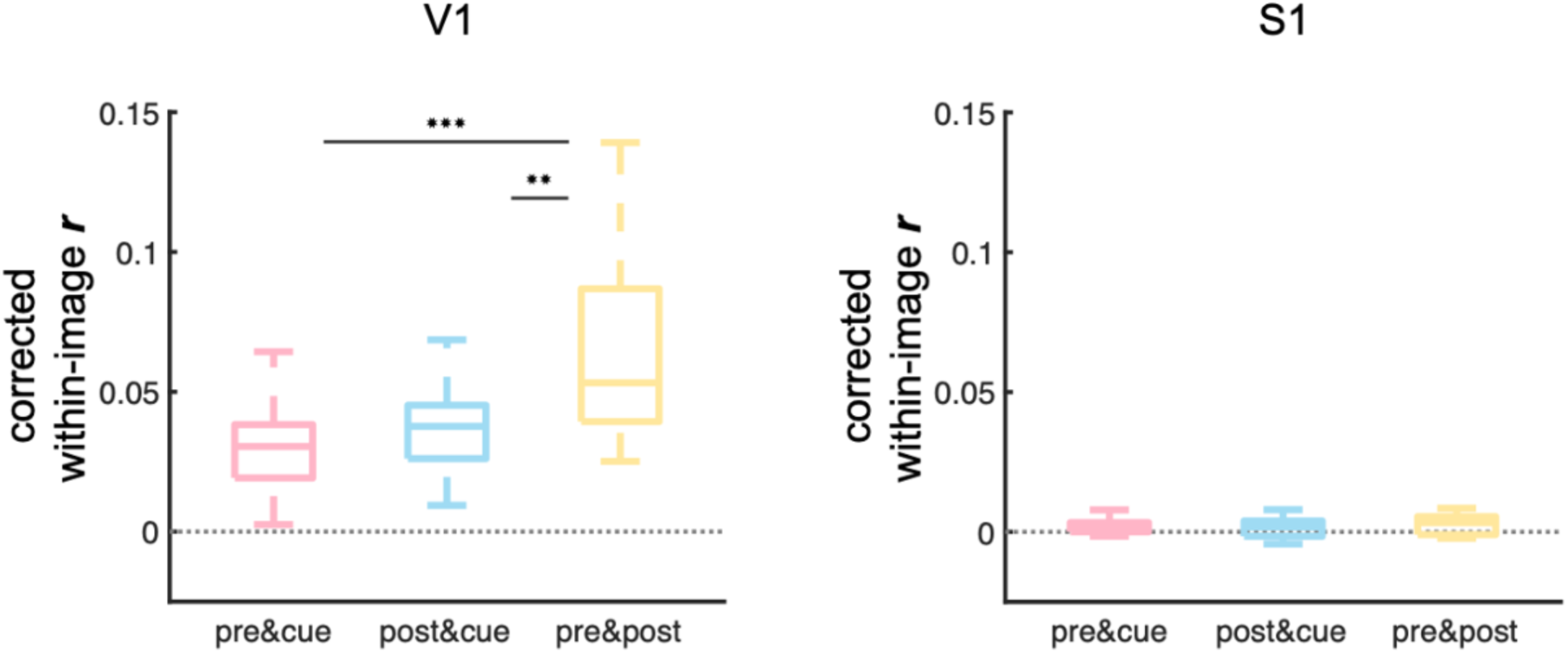
Group-level results reveal input-relevant representations in V1. Corrected mean correlations for across-condition comparisons of corresponding images in V1 (left) and S1 (right) for all subjects across all paradigms. For each across-condition comparison, within-image similarities are corrected by subtracting mean similarities of temporal controls within each image set and ROI at the single-subject level. Paired t-tests of mean corrected within-image ***r*** across all subjects were run between condition comparisons for each ROI, * denotes significance of *p* < 0.05, ** denotes p < 0.01, and *** denotes *p* < 0.001.

## Discussion

Perceptual learning, where new information alters neural representations, is widely studied with multivariate fMRI. Here we demonstrate how well-known temporal order effects can mimic such learning in the brain, by biasing representational similarity analysis measures (RSA; Kriegeskorte, 2008), revealing a critical vulnerability. Temporal artefacts, if not properly addressed, can masquerade as genuine perceptual effects, yet appropriate controls are rarely reported. This raises the possibility that some findings attributed to perceptual learning may be inflated or artefactual. To address this, we propose three methodological safeguards for multivariate fMRI studies of learning: (1) minimising order confounds, (2) controlling for temporal artefacts, and (3) checking that measured effects are specific to cognitive processes.

We tested four fMRI cueing paradigms, including ones widely used in the literature, designed to elicit perceptual reorganisation - the recognition of a previously ambiguous two-tone version of a photo image after cueing with the original photo. All paradigms featured the same basic structure: naively viewed two-tone images (‘Pre’), followed by their corresponding photo cues (‘Cue’), and the same two-tones viewed again after cueing (‘Post’). Our key manipulation was how these conditions were distributed across fMRI runs, which in turn strongly shaped the results.

Several previous studies have reported striking perceptual reorganisation of two-tone image representations in early visual cortex by prior knowledge (Flounders et al., 2019; González-García et al., 2018; Hsieh et al., 2010; van Loon et al., 2016), but have not systematically accounted for temporal order, and other non-specific, effects. Using conventional RSA approaches, we found large and significant perceptual reorganisation markers in single-subject brain responses in V1 for some designs, which were smaller or absent in others, despite each paradigm inducing a strong behavioural effect. Strikingly similar patterns appeared in a control region, the primary somatosensory cortex (S1), which is not usually involved in visual scene parsing. A genuine neural signature of perceptual reorganisation should (a) replicate across fMRI paradigms that elicit strong behavioural effects - notwithstanding potential differences in design efficiency, and (b) be largely concentrated in visual areas. As neither criteria were met in our data, we suspected a methodological rather than cognitive driver of the perceptual reorganisation results in V1.

### A three-step framework for minimising temporal confounds in RSA studies on learning

It is well known that fMRI is subject to multiple artefacts in the temporal domain (Bianciardi et al., 2009; Mumford et al., 2014; Power et al., 2012; Zarahn et al., 1997). Recent studies have reiterated the need to apply controls that fully account for temporal artefacts, such as trial randomisation (Alink et al., 2015; Cai et al., 2019; Mumford et al., 2014), or corrective analyses (Cai et al., 2016; Lee & Rideaux, 2025). However, as perceptual learning paradigms require fixed condition orders, only partial randomisation is possible. Multiple temporal order confounds are likely to affect a single dataset simultaneously (e.g., drift in multi-voxel pattern correlations within-vs across-run, over time within a run, and over the entire session), and their severity may vary from study to study depending on the scanner, the sequence, the design, and data processing steps. Given these manifold causes, a blanket recommendation for specific designs (e.g., across-run designs) that minimise a particular temporal correlation structure confound (e.g., within-run correlation structure) is unlikely to be effective. We therefore propose a targeted, flexible framework consisting of three key steps:

1. **Minimise stimulus order confounds.** As a first step, we recommend visualising the temporal correlation structure of fMRI RSA data for any study permitting only partial trial randomisation, to ensure that the contrasts of interest in the paradigm design are minimally confounded with temporal artefacts. As we show in Figure 1C, partial randomisation - for example randomising image order but not condition - can obscure temporal confounds without removing them, as RSMs are conventionally reordered by image identity (image-order RSMs) and this randomly scatters temporally ordered correlation structure across the RSM. Moreover, because non-corresponding image pairs may have different temporal placements within the session for different participants (i.e., two images presented in the same run for subject A, may appear in different runs for subject B), any confounds driven by when non-corresponding images are shown may be reduced when averaging image-ordered RSMs across subjects. In contrast, matching images will always appear in a consistent temporal relationship, so temporal order biases for these images will remain, even at the group level. In our data, visualising RSA matrices by actual presentation order in small pilot samples revealed strong temporal correlation structures at the individual participant level that were present in all paradigms and hidden in conventional analyses. Crucially, this made clear how design choices interacted with these patterns. In our data, when placing key condition comparisons within the same fMRI run, the inherent correlation structure of the BOLD measure, which was consistently more correlated for within-than across-run image pairs, produced a clear artifactual pattern resembling perceptual reorganisation (Figures 2B and 3B). Thus, run-ordering RSA matrices for a small number of pilot scans can offer critical information for design optimisation.
2. **Control for temporal order confounds.** Even after avoiding the most pronounced temporal confounds at the design stage, more subtle temporal correlation structure may still bias RSA results. It is therefore essential to apply temporally matched control comparisons during the analysis stage to isolate genuine learning-related changes. In our study of perceptual reorganisation, we corrected for temporal order effects by subtracting from the correlation of each corresponding image pair (Pre vs Cue vs Post) the mean correlation between non-matching control image pairs across these same conditions, presented at equivalent time points. These temporal controls, shown in grey in Figures 2 and 3B, shared the same temporal correlation structure confounds as the main comparison but lacked image-specific perceptual learning effects. This control provides a benchmark: for representational similarity changes to reflect true perceptual reorganisation specific to the learnt image, similarity between corresponding images needs to exceed that of the temporally matched, non-corresponding pairs. Applying this correction removed artefactual effects that mimicked perceptual reorganisation in both primary visual cortex (V1) and a control region (S1), demonstrating that uncorrected RSA measures can be misleading, in our case particularly in within-run designs. Notably, after this correction, multi-voxel patterns in V1, but not non-visual region S1, were most similar for repeated presentations of identical two-tone images, in line with low-level visual feature encoding in this region.
3. **Check for process specificity using complementary measures.** Finally, whilst effective, baseline correction alone may not fully eradicate effects of non-interest from fMRI analyses. This is because multiple cognitive and non-cognitive processes contribute to the fMRI signal at any one time, and these may interact in non-additive ways. We therefore recommend incorporating complementary measurements as negative controls to ensure that the effects measured are perceptually relevant. This could consist of a control ROI outside the pathways likely to be affected by the cognitive effect in question, as we apply here with S1. Additionally, behavioural measures of perception collected during scanning can be used to separate out trials where perceptual reorganisation effects should be weak or absent, such as two-tones for which recognition was unaltered by the photo cue (either the cue was not needed to recognise the two-tone, or the cue was ineffective), so little learning should have taken place. Notably, the control analysis in point 2 also functions as a negative control: by comparing multi-voxel pattern correlations for corresponding images to non-corresponding images, it accounts not only for temporal order effects but also for non-specific cognitive influences, such as attention or response suppression, that may apply generally across the stimuli set.

These recommendations not only help avoid temporal-order-related confounds but also ensure that observed changes truly reflect the learning process. Here, we focused on how learning affects responses to specific individual images. In contrast, other two-tone studies have looked at broader, image-identity independent changes. For example, they have shown that representations of cued two-tone images become more distinct from each other than for uncued two-tone images (González-García et al., 2018; van Loon et al., 2016), that cued two-tones become more similar to photo cues as shown using SVM photo vs two-tone classification (van Loon et al., 2016), or that a cued two-tone becomes more similar to its corresponding photo cue without ruling out that its similarity to a non-corresponding photo is also increased (Hsieh et al., 2010). All of these effects can be influenced by unrelated factors like attentional drift, repetition suppression, or adaptation (Alink et al., 2015; Cai et al., 2019; Grill-Spector et al., 2006), which over time may reduce within-condition correlations or increase apparent cue similarity without reflecting true learning. In contrast, image-specific contrasts and converging measures, as recommended here, better isolate content-driven perceptual changes, especially when combined with appropriate controls for global trends.

A limitation of our study is the small sample size of participants scanned with individual perceptual learning designs, which restricts our ability to meaningfully compare confounds across paradigms. However, the point we aim to make is principled and pragmatic, not specific to the tested task or paradigms. Our data clearly reveals how temporal correlation structure confounds present in all paradigms can be more harmful for some designs than others, and we demonstrate how even small N designs that are feasible to run as pilots can assess these inherent confounds in the data, and be used to develop protocols that minimise and control for their unwarranted impact on conclusions.

After applying the above recommended corrective steps, we no longer observed measurable perceptual reorganisation in V1; instead, V1 representations of cued two-tone images were most similar to when these two tones were first presented, and poorly recognised, consistent with low-level visual encoding of image features. Whether perceptual reorganisation of these stimuli is present at the level of V1 remains an open question, which this manuscript does not attempt to answer. While it has been reported in multiple studies so far (Flounders et al., 2019; González-García et al., 2018; van Loon et al., 2016), even to the level that perceptual reorganisation overrides bottom-up representations of the original two-tone in V1, it is unclear to what degree these results are affected by the confounds presented here. In our data, evidence of perceptual reorganisation in V1 and S1 was fully eliminated after applying appropriate corrections. We therefore recommend that all studies on neural mechanisms of learning - and perceptual reorganisation in particular - follow steps to minimise, correct, and check for non-specific effects (temporal and cognitive) that can confound fMRI measures of learning over time. Systematically accounting for temporal confounds in this way will enable accurate measures of genuine cognitive effects, and improve comparability and replicability across studies.

## Methods

### Participants

Fifteen healthy adults participated in the study. All participants had normal or corrected-to-normal vision and provided written informed consent before participating. Three participants were excluded; two due to poor task compliance and another due to a technical issue. Of the remaining twelve participants, four were female, one was left-handed, and the mean age was 25.8 years (age range: 19.8 - 30.9 years). The research was carried out in accordance with the tenets of the Declaration of Helsinki and was approved by the UCL Ethics Committee (#22397/001).

### Stimuli

Stimuli were generated from 20 images of animals and objects from The Berkeley Segmentation Dataset and Benchmark (Martin et al., 2001) using Matlab R2022a (2022). Images were converted to greyscale and cropped to a square to create photo cues. Two-tone stimuli were created using methods previously described elsewhere (Milne et al., 2024). Specifically: photo cues were smoothed with a 2D Gaussian filter and then thresholded to binarise pixel luminance to black or white, creating images with obscured edges. Smoothing levels and binarisation thresholds varied per image; standard deviation of Gaussian distribution ranged from 0.5 - 6.0 SD, binarisation thresholds ranged from 0.4 - 2.0 times the mean luminance of the greyscale image. An additional four two-tones generated with no smoothing were used in a practice task to test task comprehension. Five ‘Abstract’ two-tone images were also created by scrambling two-tone images, producing contentless two-tones for which photo cues do not exist, but are not used in analyses presented in this study. All stimuli were presented as a 13 degrees of visual angle square centred on a 50% grey background, and projected onto a screen at the back of the MRI scanner, which was viewed via a mirror.

### Procedure

During an experimental trial participants viewed a two-tone or a greyscale image and judged whether the presented image was of an animal or a man-made object. Verbal recognition of the specific item in the image was recorded after each scan in a verbal recognition task.

Each of the twelve participants completed one of four different task paradigms (see Figure 1B): Within (n = 4), Mixed-Within (n = 2), Mixed-Across (n = 3) and Across (n = 3). All paradigms included 20 images shown in three conditions: ‘Pre’ two-tone condition, ‘Cue’ greyscale photo condition and ‘Post’ two-tone condition. In each paradigm, images and conditions were distributed across fMRI runs in different ways (see below). The order of presented images was shuffled per participant, to allow for maximal randomisation while maintaining the sequential constraints of the task (see Figure 1B). Between fMRI runs, participants completed a verbal recognition task of the two-tone images to measure their pre- and post-cueing recognition.

The single-trial structure of all fMRI paradigms was identical. Each trial started with a black fixation cross presented for 1 s, followed by a 0.2 s stimulus presentation of either a photo cue or a two-tone image. This was followed by a grey fixation cross presented for 1.8 s, prompting the participant to respond via button press to whether the image was of an animal or an object. Each trial was followed by a variable inter-stimulus interval of 0.1 to 4.1 s, consisting of a grey fixation cross. In the verbal recognition task, participants saw each image for 0.2 s, and verbally answered the question ‘Can you name what you saw?’. Participants’ answers were typed on screen by the experimenter, and corrected by the participant where necessary.

Before the fMRI experiment, participants first received task instructions and completed a short practice task outside of the scanner. This task comprised eight trials of four pairs of unsmoothed two-tones and corresponding photo cues, and was repeated until the participant correctly categorised all trials as either an animal or an object.

In the Within paradigm, each fMRI run included all three conditions of four images - first as a two-tone presented before cueing (‘Pre’ condition), second as a photo cue (‘Cue’ condition), and third as a two-tone presented after cueing (‘Post’ condition). In addition, one abstract two-tone image was shown alongside Pre and Post conditions for each run, but not as a photo cue. Each image was repeated six times with a total of 84 trials per run. Presentation order was pseudorandomised across images within the same condition, but not across conditions. To distribute the image repeats, we shuffled image orders of each condition within 3 trial blocks including 2 repeats of each image, which were pseudorandomised ensuring no image was repeated twice in a row. Three rest periods of 13 s duration were included at the start, middle (between Pre and Cue conditions) and end of each run, where a grey fixation cross was shown. In total, the paradigm included five fMRI runs that each lasted 7 m 47 s. The verbal recognition task was completed between each run for the four two-tone images shown in the previous run (post-cueing recognition) and the four two-tones to be shown in the next run (pre-cueing recognition), and at the start and end of the experiment to test pre-cueing recognition of the first four two-tones, and post-cueing recognition of the last four two-tones, respectively.

In the Mixed-Within paradigm, fMRI runs included four images presented in Pre condition, and four different images presented in both Cue and Post conditions. In addition, one abstract image was shown alongside Pre and Post conditions for each run, but not as a photo cue. Each image in Pre and Post conditions was repeated six times and images presented in Cue condition were repeated three times. In total there were 72 trials per run when all conditions were included (see below). Presentation order was pseudo-randomised across the interspersed Pre and Post conditions - which contained different images. In the Cue condition, randomisation of trial order was carried out across images but not mixed with the other conditions. Trials were divided into two types of blocks: two-tone blocks where Pre, Post, and Abstract were combined and Cue blocks. Each two-tone block included two repetitions of ten images in a pseudorandomised order. Cue blocks included only one repetition of four images in a pseudorandomised order. Both blocks were repeated 3 times, and Cue and two-tone blocks were interleaved, so a Cue block was always followed by a two-tone block. Three rest periods of 10 s duration were included at the start, middle (between two-tone and Cue conditions) and end of each run. The nature of this paradigm meant that we could not start with a run that includes all three conditions, e.g., to have images to show in the Cue and Post conditions they must first be presented in a separate run with a Pre condition. Therefore the paradigm included six fMRI runs of different lengths: run 1 included only the Pre condition (2 m 55 s), runs 2-5 included all conditions (6 m 35 s), and run 6 included only Cue and Post conditions (3 m 55 s). The verbal recognition task was completed after each run for any two-tone images shown in the previous run (pre- and post-cueing recognition).

In the Mixed-Across paradigm, each condition of corresponding images was presented in a separate run. fMRI runs included four images presented in Pre condition, four different images presented in Cue condition, and four different images presented in Post condition. In addition, one abstract image was shown alongside Pre and Post conditions for each run, but not as a photo cue. Images in Pre and Post conditions were repeated six times, and images presented in Cue condition were repeated three times. In total there were 72 trials per run when all conditions were included (see below). Because all conditions within a run were of different images, presentation order was pseudorandomised across all trials (i.e., across images and conditions). To distribute the image repeats across the run, we shuffled image orders within 3 trial blocks including 2 repeats of the 4 Pre, and 4 Post two-tones, and 1 repeat of the 4 photo cues. Three rest periods of 10 s duration were included at the start, middle (between two-tone and ‘Cue’ conditions) and end of each run, where a grey fixation cross was shown. The nature of this paradigm meant that we could not start with a run that includes all three conditions, e.g., to have images to show in the Cue condition they must first be presented in a separate run in Pre condition. Therefore the paradigm included seven fMRI runs of different lengths: run 1 included only the Pre condition (2 m 52 s), run 2 included Pre and Cue conditions (4 m 3 s), runs 3-5 included all conditions (6 m 33 s), run 6 included Cue and Post conditions (4 m 3 s), and run 7 included only Post condition (2 m 52 s). The verbal recognition task was completed after each run for any two-tone images shown in the previous run (pre- and post-cueing recognition).

In the Across paradigm, each condition (Pre, Cue or Post) was presented in a completely separate run. Each fMRI run included 10 images (half the stimuli set randomly split). In addition, two abstract images were shown in Pre and Post condition runs, but not in Cue runs. Each image was repeated six times, for a total of 72 trials per Pre/Post runs, and 60 trials per Cue run. To distribute the image repeats across the run, we shuffled image orders within 3 trial blocks including 2 repeats of each image. Three rest periods of 13 s duration were included at the start, middle and end of each run, where a grey fixation cross was shown. In total, the paradigm included two Pre condition runs (runs 1 and 4) that lasted 6 m 46 s each, two Cue condition runs (runs 2 and 5) that lasted 5 m 45 s each, and two Post condition runs (runs 3 and 6) that lasted 6 m 46 s each. The verbal recognition task was completed after each run that included two-tone images (runs 1, 3, 4, and 6) for the ten Pre or Post condition two-tone images shown in the previous run.

### Data acquisition and preprocessing

Data were collected at the Birkbeck-UCL Centre for NeuroImaging (BUCNI, London, UK), on a Siemens PRISMA 3 Tesla scanner and a 32-channel head coil (identical to 32-channel but without view-obstructing front). T2*-weighted echo-planar imaging were collected with an accelerated multi-band sequence by CMRR (version R016a, https://www.cmrr.umn.edu/multiband; Cauley et al., 2014; Xu et al., 2013); multi-band factor: 4, voxel resolution: 2mm isotropic, FOV: 212×212×96mm, flip angle: 60°, repetition time (TR): 1000ms, echo time (TE): 35.2ms, echo spacing: 0.56ms, bandwidth: 2620Hz/Px, with 48 transverse slices angled to be approximately parallel to the calcarine sulcus whilst avoiding the orbital cavities).

A T1-weighted structural image was acquired with a 32-channel coil, using MPRAGE sequence (voxel resolution: 1mm isotropic, 208 slices, FOV: 256 x 256 x 208mm, flip angle: 9°, TR = 2.3 s, TE = 2.98 ms, TI = 900 ms, bandwidth: 240Hz/Px, acquisition time: 5min30s). Preprocessing of structural and functional data was performed with FSL (FMRIB’s Software Library, www.fmrib.ox.ac.uk/fsl). Structural images were skull-stripped, and functional images were motion assessed using framewise displacement (threshold = 0.9). Functional data processing was carried out using FSL’s FEAT (FMRI Expert Analysis Tool) Version 6.00 (www.fmrib.ox.ac.uk/fsl), including: motion correction using MCFLIRT (Jenkinson et al., 2002), non-brain removal using BET (Smith, 2002); spatial smoothing using a Gaussian kernel of FWHM 3 mm, grand-mean intensity normalisation by a single multiplicative factor, high-pass temporal filtering (Gaussian-weighted least-squares straight line fitting, with sigma=50.0s). Time-series statistical analysis was carried out using FILM with local autocorrelation correction (Woolrich et al., 2001). Specifically, a general linear model (GLM) was constructed at the single-subject level in native functional space. Regressors for each image (Im_1_ to Im_20_) in each condition (Pre, Cue and Post) that included all trial repeats of that stimulus, as well as regressors of suprathreshold movement confounds, were convolved with the hemodynamic response function (HRF) following a double gamma function. First-order temporal derivatives of each regressor were also included in the model. Functional images were registered to high resolution structural scans using FLIRT (Jenkinson et al., 2002; Jenkinson & Smith, 2001) with boundary-based registration (BBR), and these high resolution images were registered to a 2 x 2 x 2 mm MNI atlas using FNIRT nonlinear registration (Andersson et al., 2007a, 2007b) with 12 degrees of freedom.

### Region of interest (ROI) definition

ROIs of primary visual cortex (V1) and primary somatosensory cortex (S1) were anatomically defined for each participant. For V1, cortical reconstruction was performed in FreeSurfer 7.3.2 (https://surfer.nmr.mgh.harvard.edu/fswiki/recon-all). A cortical V1 label was then obtained using Benson retinotopy (Benson & Winawer, 2018), which was restricted to an eccentricity of 9 degrees of visual angle to match the visually stimulated region. Restricted V1 labels were then converted to volumetric masks using FreeSurfer’s BBregister and combined across hemispheres. For S1, volumetric masks were anatomically defined in standard space using the Harvard-Oxford Cortical Structural Atlas (https://fsl.fmrib.ox.ac.uk/fsl/fslwiki/Atlases) template region for the Postcentral gyrus, and registered back to the native space of each subject.

### Representational similarity analysis (RSA)

For each ROI (V1 and S1) the GLM beta-weights per run were extracted and normalised with multivariate noise normalisation (Walther et al., 2016) using the Representational Similarity Analysis Toolbox (Nili et al., 2014) adapted to FSL (https://zenodo.org/records/1112437) in MATLAB version R2024b (MathWorks, 2024). Pairwise comparisons between normalised multivariate patterns were performed using Pearson correlation as the distance measure. All pairwise comparisons were then visualised in a 60×60 Representational Similarity Matrix (RSM). In the RSM, images were arranged along the rows and columns based on two factors: the condition in which the image appeared (i.e., rows 1-20 = Pre, rows 21-40 = Cue, and rows 41-60 = Post) and within each condition either (1) an arbitrary ordering of image identities (‘image-order’) or (2) the order in which the images were presented to each individual subject during scanning (‘run-order’). Specifically, the sequence by which run-order RSMs are arranged corresponds to which fMRI run a given condition of an image appeared. For example, Within, Mixed-Within and Mixed-Across paradigms present four unique images in Pre condition per run, so correlations of the Pre condition images presented in run 1 are displayed in run-order RSM rows 1-4, images presented in run 2 are displayed in rows 5-8, and so on. The same order applies across all three stimulus conditions, with subsequent conditions presented according to the paradigm design (see Figure 1B). The order of images within these four-row blocks is arbitrary, as images are repeated multiple times within a run in blocks with different orders. Ordering RSMs by this order for each stimulus condition generates a matrix structure where images presented in the same run appear in consecutive rows within each condition block, while corresponding images across conditions can be identified along the off-diagonal sections of the RSM.

Both image-order and run-order RSMs were also constructed and analysed using non-normalised beta values with similar results (see Supplementary 3).

### Temporal controls

To control for temporal artefacts in across-condition comparisons of corresponding images, we computed temporal controls using non-corresponding images that had similar temporal presentation patterns. These captured the same temporal correlation structure but excluded content-specific information. A temporal control was calculated for each set of images that were grouped together in the experimental design (sets of 4 images in Within, Mixed-Within, and Mixed-Across paradigms; sets of 10 images for Across paradigm).

In the ’Within’ paradigm, all corresponding image comparisons occur within the same run. To create temporal controls for Pre&Cue comparisons, for example, we calculated correlations between all non-corresponding Pre and Cue images within each run (e.g., for a 4-image set size, this yields 12 correlations: Pre image1 with Cue images 2, 3, & 4, Pre image2 with Cue images 1, 3, & 4, etc.). These correlations were averaged to create a run-specific temporal control value for the Pre&Cue comparison. To correct for temporal artefacts, each corresponding image correlation (e.g., Pre image1 & Cue image1) was then corrected by subtracting the average temporal control correlation value taken from its respective run. This process was repeated for all across-condition comparisons (Pre&Cue, Post&Cue, Pre&Post).

In the Mixed-Within paradigm, the temporal structure differs, but the logic is the same. In this design, Pre images from one run are compared with their corresponding Cue and Post images from the subsequent run. For Pre&Cue temporal controls, we therefore calculated correlations between non-corresponding Pre and Cue images that maintained this across-run structure (e.g., Pre image1 in run 1 with Cue images 2, 3, & 4 in run 2). Similarly, Pre&Post temporal controls used non-corresponding images spanning these different runs. However, Post&Cue comparisons had a different structure: both Post and Cue images appeared within the same run, so temporal controls were calculated between non-corresponding Post and Cue images within that run (e.g., Post image1 with Cue images 2, 3, & 4, all in run 2).

In the Mixed-Across paradigm, all comparisons span different runs (e.g., Pre in run 1, Cue in run 2, Post in run 3), so temporal controls matched these specific across-run distances. For Pre&Cue comparisons, temporal controls were calculated using correlations between non-corresponding Pre and Cue images that maintained the same run separation (e.g., for a 4-image set, this yields 12 correlations: Pre image 1 in run 1 with Cue images 2, 3, & 4 in run 2, etc.). These correlations were averaged to create temporal control values that captured the across-run correlation structure. The same logic applied to Post&Cue and Pre&Post comparisons, with temporal controls always matching the specific run distances of the corresponding image comparisons.

The Across paradigm had a fundamentally different structure: each stimulus condition was presented in dedicated runs (runs 1 and 4 for Pre, runs 2 and 5 for Cue, runs 3 and 6 for Post). Additionally, the Across paradigm used larger image sets (10 images per set instead of 4), resulting in 90 non-corresponding correlations averaged for each run-specific temporal control (10×10 matrix excluding 10 diagonal pairs). To correct for temporal artefacts, for example, the correlation between Pre image 1 & Cue image 1 (where Pre image 1 was presented in run 1 and Cue image 1 in run 2) would be corrected by subtracting the temporal control value calculated from all non-corresponding correlations between Pre images in run 1 and Cue images in run 2 (e.g., Pre image 1 & Cue image 2, Pre image 2 & Cue image1, etc.).

### Statistical analyses

All statistical analyses were carried out in Matlab version R2024b, equipped with the Statistics and Machine Learning Toolbox and the Bioinformatics Toolbox (MathWorks Inc., 2024).

To assess perceptual reorganisation behaviourally, we modelled subjects’ behavioural responses to the image recognition and image categorisation tasks they performed. For each paradigm, all trials for all participants were analysed using a generalised linear mixed model (function fitglme) with random intercepts for subjects and images on accuracy of image recognition responses (a binomial measure of image recognition recorded as either recognised or unrecognised once for each image in Pre and Post conditions), and a linear mixed model (function fitlme) with the same random intercepts on accuracy of image categorisation responses (for each image, an average of the six repeated image categorisation scores in Pre and Post conditions).

To assess neural perceptual reorganisation in each cortical ROI (V1 and S1), we calculated pairwise correlations between representations of corresponding images (within-image ***r***) in each stimulus condition (Pre&Cue, Post&Cue and Pre&Post) at the single-subject level. We then ran paired t-tests between the three across-condition comparisons of all 20 image pairs within each subject and ROI. Multiple comparison corrections were performed using FDR (see Supplementary 4).

To test for differences between within- and between-run correlations across all paradigms, we ran a linear mixed-effects model (function fitlme) of Fisher z-transformed across-condition correlations of V1 responses to corresponding images with random intercepts for subjects and images, testing for main effects of comparison type (within- or between-run) and condition pair (Pre&Cue, Post&Cue or Pre&Post), as well as the interaction between these factors.

To test for differences in representational similarity across the conditions due to temporal correlation structure unrelated to stimulus identity, we ran paired t-tests between the three across-condition comparisons of all temporal controls, and ran FDR multiple comparisons corrections (see Supplementary 5).

To test for the isolated effects of temporal artefacts, and the underlying perceptual reorganisation effects after these have been removed, we ran paired t-tests between each across-condition comparison (Pre&Cue, Post&Cue, and Pre&Post) of all 20 image pairs within each subject and ROI for corrected within-image similarities, and ran FDR multiple comparisons corrections (see Supplementary 6). We further combined data from participants across all paradigms and ran group-level paired t-tests on corrected V1 and S1 within-image similarities to assess for neural markers of perceptual reorganisation (Figure 4).

## Data and Code Availability

All analysis code will be publicly available following publication.

## Author Contributions

**Georgia Milne:** Conceptualisation, Investigation, Methodology, Formal analysis, Writing - Original Draft, Writing - Review & Editing, Visualisation; **Hugo Chow-Wing-Bom:** Investigation; **Kim Staeubli:** Investigation; **John Greenwood:** Writing - Review & Editing; **Peter Kok:** Writing - Review & Editing; **Roni Maimon Mor:** Supervision, Conceptualisation, Methodology, Writing - Original Draft, Writing - Review & Editing; **Tessa Dekker:** Supervision, Conceptualisation, Methodology, Writing - Original Draft, Writing - Review & Editing.

