## Supplementary Information for "Temporal confounds emulate multivariate fMRI measures of perceptual learning"

### Supplementary 1: Behavioural perceptual reorganisation effect

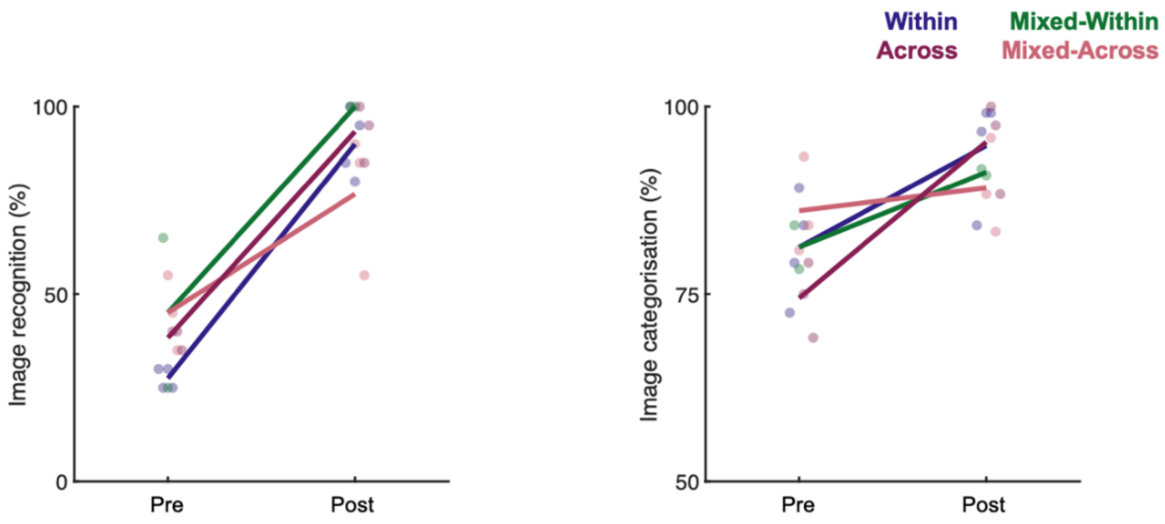


Photo cueing improved recognition of two-tones in both behavioural measures; verbal naming of image content completed between fMRI runs (‘image recognition’, left plot) and button-press animate/inanimate categorisation completed within each trial (‘image categorisation, right plot). Lines show mean cueing effect per paradigm (mean Post recognition - mean Pre recognition), markers show individual subject performance. A generalised linear mixed model with random intercepts for subjects and images revealed a significant effect of two-tone condition on image recognition for each paradigm ($\beta > 0.31$, SE $< 0.08$, $t > 4.65$, $p < 10^{-5}$), and a linear mixed model with the same random intercepts revealed a significant effect of two-tone condition on image categorisation for Within, Mixed-Across and Across paradigms ($\beta > 0.1$, SE $< 0.047$, $t > 3.7$, $p < 0.026$), and a nonsignificant trend in the Mixed-Within paradigm ($\beta = 0.11$, SE $= 0.06$, $t = 1.67$, $p = 0.098$).

### Supplementary 2: Image- and run-order single-subject RSMs

Temporal order artefacts are substantial at the single-subject level, causing large amplifications of within-image similarities for any comparison made within-run (Supplementary Figure 2). These paradigm-dependent patterns are highly consistent at the single-subject level; Within each paradigm, pairwise pearson correlations between subjects' run-order RSM's were very high: V1 = 0.92, S1=0.94 (averaged across all paradigms), and significantly larger than image-order RSMs: V1 = 0.61, S1 = 0.60 (paired t-test between run-order and image-order pairwise correlations across all paradigms, for each ROI: $t(12) > 7.5$, Cohen's $dz > 2$, $p < 10^{-5}$).

**Supplementary Figure 2: Representative single-subject representational similarity matrices in V1 and S1, image-order & run order RSMs**

**A** Representative single-subject image-order RSMs from each paradigm group shown for primary visual cortex (V1), ordered by stimulus condition (Pre, Cue and Post) and image identity ($Im_1$ to $Im_{20}$). **B** Representative single-subject run-order RSMs from each paradigm group for V1, ordered by stimulus condition (Pre, Cue and Post) and experiment presentation order. Subjects are the same as in panel A. **C** Representative single-subject image-order RSMs from each paradigm group shown for primary somatosensory cortex (S1). Subjects are the same as in panel A. **D** Representative single-subject run-order RSMs from each paradigm group for S1. Subjects are the same as in panel A.

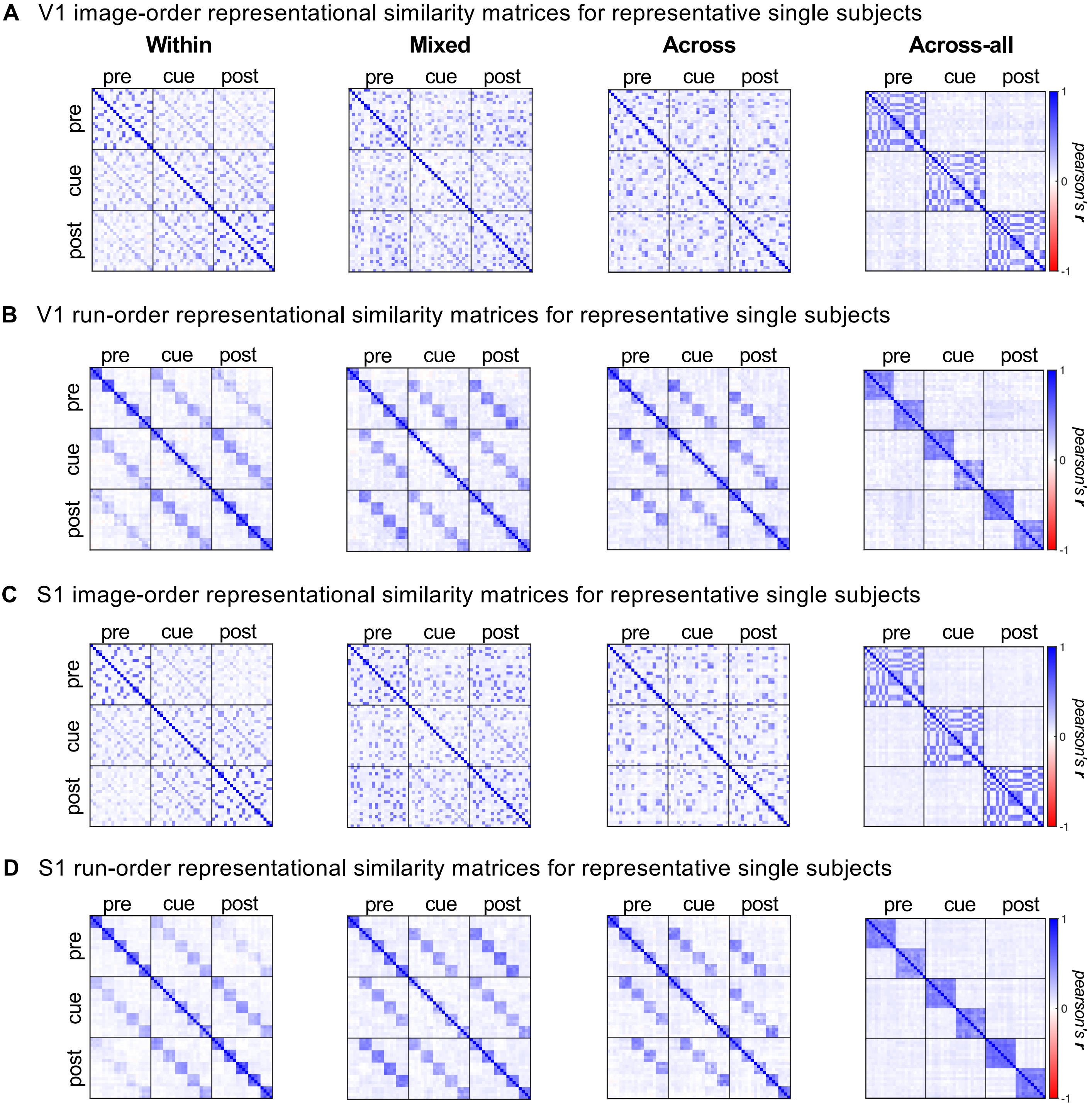
A V1 image-order representational similarity matrices for representative single subjects
Within
Mixed
Across
Across-all
pre
cue
post
pearson's r
1
0
-1
B V1 run-order representational similarity matrices for representative single subjects
C S1 image-order representational similarity matrices for representative single subjects
D S1 run-order representational similarity matrices for representative single subjects

### Supplementary 3 - Unnormalised RSA measures of perceptual reorganisation

We applied a widely used multivariate normalisation of fMRI pattern data (Walther et al. 2016), which increases the SNR of our data, by removing correlated noise from the patterns, increasing signal in similarity matrices. After this normalisation the temporal artefacts became very apparent in our data. It is important to note, however, that the artefacts were not *introduced* by the multivariate normalisation step – below we present evidence for that they become more detectible after normalisation because this procedure (a) increases SNR, and therefore power to detect the impacts of temporal order effects that (b) were already present in unnormalised data.

Firstly, as expected with lower SNR data, pair-wise correlations in the unnormalised analysis of V1 were on average higher and more variable across all subjects (increased mean and standard deviation in unnormalised compared to normalised $p < 0.01$). Further evidence for lower SNR in the unnormalised data can be seen when comparing corresponding image correlations (e.g. Pre image1 with Post image1) to correlations across noncorresponding images (e.g. Pre image1 and Post image2, also referred to as temporal controls). Across all paradigms, these were only significant in 4 of the 12 participants in the unnormalised data analysis. After applying multivariate normalisation these became significant in 11 out of 12 ($p < 0.05$) and trending in one ($p = 0.067$). Thus multivariate normalisation increases SNR, rendering measures more sensitive to signals due to stimulus manipulations as well as temporal correlation structure that varies across the timeseries, which is not controlled for by this analysis.

Secondly, temporal order artefacts are already present in our unnormalized data (Supplementary Figure 3). For example, comparing across un-normalised data from all paradigms, correlations of corresponding images across conditions presented within the same run (within-run) were higher than those presented in different runs (between-run; linear mixed model: $\beta$(within/between) = 0.12, SE = 0.049, $t(714) = 2.51$, $p < 0.05$, 95% CI [0.027, 0.217]). When we perform the main analyses steps with unnormalised data we observe similar artefactually enhanced effects of perceptual reorganisation (Post&Cue > Pre&Cue) for comparisons of corresponding images, as well as in the temporal controls (Supplementary Figure 3A and 3B, 2nd rows). Although, while these patterns are numerically similar as in the main analyses they did not always reach significance at the single-subject level, likely due to lower SNR in the unnormalised data.

The unnormalised data analysis showed similar artefactually enhanced effects of perceptual reorganisation (Post&Cue > Pre&Cue) in the across-condition comparisons of corresponding images, as well as in the temporal controls (Supplementary Figure 3A and 3B, 2nd rows). However, while these are numerically similar they did not always reach significance at the single-subject level, likely due to lower SNR in the unnormalised data as described above. In the Mixed-Within paradigm, significant effects were present for all single-subject comparisons of temporal controls in S1 and V1 ($p < 0.05$), and all comparisons of corresponding images ($p < 0.01$) except one ($p = 0.16$). In the Within paradigm, Post&Cue > Pre&Cue was significant in V1 temporal controls for two of the three subjects that showed a significant effect in the normalised data ($p < 0.05$) and trending for one ($p = 0.08$). These were also present in corresponding images for one subject in V1 ($p = 0.011$) and three subjects in S1 ($p < 0.05$). In the Mixed-Across paradigm, no significant effects of perceptual reorganisation were present for corresponding images or temporal confounds in either V1 or S1 ($p > 0.05$ for all subjects), whilst for the Across paradigm, these effects reached significance in at least one ROI for one subject of three subjects ($p < 0.05$). Together, this suggests the temporal artefacts observed in normalised data are also present at the single-subject level before applying normalisation.

After correcting for temporal order artefacts (see Methods), no significant perceptual reorganisation effects were found at the single-subject level for any paradigm in either ROI ($p > 0.05$ for all subjects). As with the normalised data, group-level comparisons of the corrected correlations (across all paradigms) revealed significant effects of Pre&Post > Pre&Cue ($p < 0.01$) and Pre&Post > Post&Cue ($p < 10^{-4}$) in V1, but no significant effects of perceptual reorganisation in either ROI ($p > 0.05$).

**Supplementary Figure 3: Unnormalised RSMs and measures of perceptual reorganisation**

**A** First row: Group-averaged run-order RSMs of unnormalised multi-voxel response patterns for primary visual cortex (V1), ordered by stimulus condition (Pre, Cue and Post) and experiment presentation order. 2nd row: Mean within-image similarities for across-condition comparisons of unnormalised multi-voxel response patterns in V1. Error bars show bootstrapped 95% confidence intervals for individual subjects, horizontal coloured bars show group averages for each across-condition comparison. 3rd row: Mean similarities of temporal controls for across-condition comparisons of unnormalised multi-voxel response patterns in V1. Error bars show bootstrapped 95% confidence intervals for individual subjects, horizontal grey bars show group averages for each across-condition comparison. 4th row: Correction of mean similarities for across-condition comparisons of unnormalised multi-voxel response patterns to corresponding images in V1. For each across-condition comparison, within-image similarities are corrected by subtracting mean similarities of temporal controls within each image set and ROI at the single-subject level. Error bars show bootstrapped 95% confidence intervals for individual subjects, horizontal hollow bars show group averages for each across-condition comparison. **B** As in panel A, but for unnormalised multi-voxel response patterns for primary somatosensory cortex (S1).

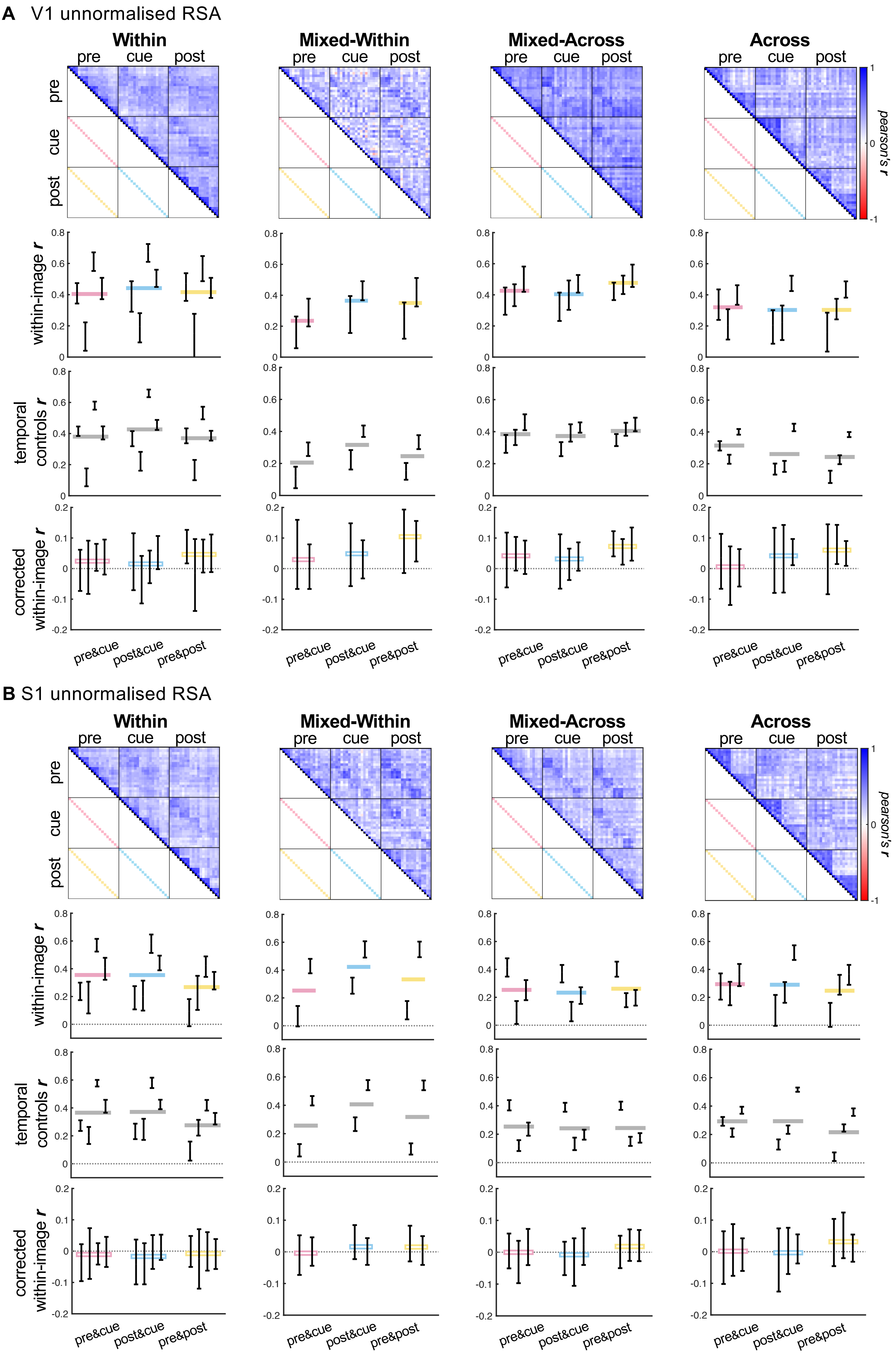

A V1 unnormalised RSA
Within
Mixed-Within
Mixed-Across
Across
pre
cue
post
pearson's r
within-image r
temporal controls r
corrected within-image r
pre&cue
post&cue
pre&post
B S1 unnormalised RSA

### Supplementary 4: Single-subject statistical tests for mean representational similarity of within-image comparisons

| V1 | Pre&Cue vs. Post&Cue | Post&Cue vs. Pre&Post | Pre&Cue vs. Pre&Post |
|---|---|---|---|
| **Within** | | | |
| subject 1 | ***p* = 0.019, *q* = 0.034, t(19) = 2.6, Cohen's *dz* = 0.57, SD: 0.071** | ***p* = 0.0091, *q* = 0.018, t(19) = -2.9, Cohen's *dz* = -0.65, SD: 0.066** | *p* = 0.91, *q* = 0.94, t(19) = -0.11, Cohen's *dz* = -0.025, SD: 0.089 |
| subject 2 | ***p* = 0.02, *q* = 0.036, t(19) = -2.5, Cohen's *dz* = -0.57, SD: 0.14** | ***p* = 0.0017, *q* = 0.0042, t(19) = 3.6, Cohen's *dz* = 0.82, SD: 0.13** | *p* = 0.076, *q* = 0.12, t(19) = 1.9, Cohen's *dz* = 0.42, SD: 0.07 |
| subject 3 | ***p* = 1.9e-06, *q* = 9.3e-06, t(19) = -6.7, Cohen's *dz* = -1.5, SD: 0.057** | ***p* = 1.8e-09, *q* = 1.6e-08, t(19) = 11, Cohen's *dz* = 2.4, SD: 0.061** | ***p* = 1e-04, *q* = 4.2e-04, t(19) = 4.9, Cohen's *dz* = 1.1, SD: 0.055** |
| subject 4 | ***p* = 0.002, *q* = 0.0045, t(19) = -3.6, Cohen's *dz* = -0.8, SD: 0.085** | ***p* = 0.016, *q* = 0.03, t(19) = 2.6, Cohen's *dz* = 0.59, SD: 0.087** | *p* = 0.22, *q* = 0.31, t(19) = -1.3, Cohen's *dz* = -0.28, SD: 0.058 |
| **Mixed-Within** | | | |
| subject 5 | ***p* = 2.9e-10, *q* = 3.5e-09, t(19) = -12, Cohen's *dz* = -2.7, SD: 0.1** | ***p* = 5.9e-09, *q* = 4.2e-08, t(19) = 9.9, Cohen's *dz* = 2.2, SD: 0.089** | ***p* = 4.9e-08, *q* = 3e-07, t(19) = -8.7, Cohen's *dz* = -1.9, SD: 0.035** |
| subject 6 | ***p* = 3.5e-17, *q* = 2e-15, t(19) = -29, Cohen's *dz* = -6.5, SD: 0.046** | ***p* = 6.1e-14, *q* = 1.5e-12, t(19) = 19, Cohen's *dz* = 4.3, SD: 0.058** | ***p* = 1.3e-05, *q* = 5.8e-05, t(19) = -5.8, Cohen's *dz* = -1.3, SD: 0.039** |
| **Mixed-Across** | | | |
| subject 7 | *p* = 0.23, *q* = 0.31, t(19) = -1.3, Cohen's *dz* = -0.28, SD: 0.038 | ***p* = 0.024, *q* = 0.041, t(19) = -2.5, Cohen's *dz* = -0.55, SD: 0.043** | ***p* = 1.3e-04, *q* = 4.7e-04, t(19) = -4.8, Cohen's *dz* = -1.1, SD: 0.032** |
| subject 8 | *p* = 0.51, *q* = 0.62, t(19) = -0.67, Cohen's *dz* = -0.15, SD: 0.05 | ***p* = 0.0043, *q* = 0.0092, t(19) = -3.2, Cohen's *dz* = -0.72, SD: 0.038** | ***p* = 0.0019, *q* = 0.0045, t(19) = -3.6, Cohen's *dz* = -0.8, SD: 0.043** |
| subject 9 | *p* = 0.57, *q* = 0.67, t(19) = -0.58, Cohen's *dz* = -0.13, SD: 0.036 | ***p* = 4.4e-04, *q* = 0.0012, t(19) = -4.2, Cohen's *dz* = -0.95, SD: 0.043** | ***p* = 2.9e-04, *q* = 8.8e-04, t(19) = -4.4, Cohen's *dz* = -0.99, SD: 0.046** |
| **Across** | | | |
| subject 10 | *p* = 0.98, *q* = 0.98, t(19) = -0.026, Cohen's *dz* = -0.0057, SD: 0.041 | *p* = 0.78, *q* = 0.86, t(19) = 0.29, Cohen's *dz* = 0.064, SD: 0.036 | *p* = 0.85, *q* = 0.9, t(19) = 0.19, Cohen's *dz* = 0.043, SD: 0.048 |
| subject 11 | *p* = 0.82, *q* = 0.88, t(19) = -0.23, Cohen's *dz* = -0.051, SD: 0.043 | ***p* = 2.6e-04, *q* = 8.1e-04, t(19) = -4.5, Cohen's *dz* = -1, SD: 0.038** | ***p* = 0.0011, *q* = 0.0028, t(19) = -3.8, Cohen's *dz* = -0.86, SD: 0.047** |
| subject 12 | *p* = 0.48, *q* = 0.6, t(19) = 0.71, Cohen's *dz* = 0.16, SD: 0.042 | ***p* = 2.2e-04, *q* = 7.4e-04, t(19) = -4.5, Cohen's *dz* = -1, SD: 0.029** | ***p* 0.0094, *q* = 0.018, t(19) = -2.9, Cohen's *dz* = -0.65, SD: 0.035** |

| S1 | Pre&Cue vs. Post&Cue | Post&Cue vs. Pre&Post | Pre&Cue vs. Pre&Post |
|---|---|---|---|
| **Within** | | | |
| subject 1 | *p* = 0.78, *q* = 0.86, t(19) = 0.28, Cohen's *dz* = 0.064, SD: 0.089 | ***p* = 0.0087, *q* = 0.018, t(19) = 2.9, Cohen's *dz* = 0.65, SD: 0.09** | *p* = 0.035, *q* = 0.057, t(19) = 2.3, Cohen's *dz* = 0.51, SD: 0.13 |
| subject 2 | ***p* = 2.3e-04, *q* = 7.4e-04, t(19) = -4.5, Cohen's *dz* = -1, SD: 0.073** | ***p* = 9.5e-09, *q* = 6.2e-08, t(19) = 9.6, Cohen's *dz* = 2.2, SD: 0.052** | *p* = 0.033, *q* = 0.055, t(19) = 2.3, Cohen's *dz* = 0.52, SD: 0.073 |
| subject 3 | ***p* = 3.9e-04, *q* = 0.0011, t(19) = -4.3, Cohen's *dz* = -0.96, SD: 0.1** | ***p* = 2.5e-10, *q* = 3.5e-09, t(19) = 12, Cohen's *dz* = 2.7, SD: 0.078** | ***p* = 4e-07, *p* = 2.1e-06, t(19) = 7.5, Cohen's *dz* = 1.7, SD: 0.068** |
| subject 4 | ***p* = 4.2e-04, *q* = 0.0012, t(19) = -4.3, Cohen's *dz* = -0.95, SD: 0.085** | ***p* = 2.1e-07, *q* = 1.2e-06, t(19) = 7.9, Cohen's *dz* = 1.8, SD: 0.085** | ***p* = 1.9e-05, *p* = 7.8e-05, t(19) = 5.7, Cohen's *dz* = 1.3, SD: 0.054** |
| **Mixed-Within** | | | |
| subject 5 | ***p* = 2e-09, *q* = 1.6e-08, t(19) = -11, Cohen's *dz* = -2.4, SD: 0.1** | ***p* = 2e-09, *q* = 1.6e-08, t(19) = 11, Cohen's *dz* = 2.4, SD: 0.1** | *p* = 0.75, *q* = 0.86, t(19) = -0.32, Cohen's *dz* = -0.072, SD: 0.038 |
| subject 6 | ***p* = 7.4e-15, *q* = 2.7e-13, t(19) = -22, Cohen's *dz* = -4.8, SD: 0.06** | ***p* = 3.4e-13, *q* = 6.1e-12, t(19) = 18, Cohen's *dz* = 3.9, SD: 0.069** | ***p* = 0.012, *q* = 0.023, t(19) = -2.8, Cohen's *dz* = -0.62, SD: 0.028** |
| **Mixed-Across** | | | |
| subject 7 | *p* = 0.79, *q* = 0.86, t(19) = 0.27, Cohen's *dz* = 0.061, SD: 0.027 | *p* = 0.24, *q* = 0.32, t(19) = 1.2, Cohen's *dz* = 0.27, SD: 0.037 | *p* = 0.19, *q* = 0.27, t(19) = 1.4, Cohen's *dz* = 0.31, SD: 0.038 |
| subject 8 | *p* = 0.93, *q* = 0.94, t(19) = -0.087, Cohen's *dz* = -0.019, SD: 0.044 | *p* = 0.16, *q* = 0.24, t(19) = -1.4, Cohen's *dz* = -0.32, SD: 0.043 | *p* = 0.13, *q* = 0.2, t(19) = -1.6, Cohen's *dz* = -0.35, SD: 0.042 |
| subject 9 | *p* = 0.32, *q* = 0.42, t(19) = 1, Cohen's *dz* = 0.23, SD: 0.041 | ***p* = 0.0027, *q* = 0.0058, t(19) = -3.5, Cohen's *dz* = -0.77, SD: 0.041** | *p* = 0.041, *q* = 0.065, t(19) = -2.2, Cohen's *dz* = -0.49, SD: 0.045 |
| **Across** | | | |
| subject 10 | ***p* = 1.2e-04, *q* = 4.7e-04, t(19) = 4.8, Cohen's *dz* = 1.1, SD: 0.023** | *p* = 0.89, *q* = 0.92, t(19) = 0.15, Cohen's *dz* = 0.033, SD: 0.031 | ***p* = 0.002, *q* = 0.0045, t(19) = 3.6, Cohen's *dz* = 0.8, SD: 0.032** |
| subject 11 | *p* = 0.036, *q* = 0.057, t(19) = 2.3, Cohen's *dz* = 0.51, SD: 0.034 | *p* = 0.29, *q* = 0.38, t(19) = -1.1, Cohen's *dz* = -0.24, SD: 0.036 | *p* = 0.37, *q* = 0.47, t(19) = 0.92, Cohen's *dz* = 0.21, SD: 0.042 |
| subject 12 | *p* = 0.56, *q* = 0.67, t(19) = 0.6, Cohen's *dz* = 0.13, SD: 0.032 | *p* = 0.68, *q* = 0.79, t(19) = 0.41, Cohen's *dz* = 0.093, SD: 0.033 | *p* = 0.23, *q* = 0.31, t(19) = 1.2, Cohen's *dz* = 0.28, SD: 0.027 |

### Supplementary 5: Single-subject statistical tests for mean representational similarity of temporal controls

| V1 | Pre&Cue vs. Post&Cue | Post&Cue vs. Pre&Post | Pre&Cue vs. Pre&Post |
|---|---|---|---|
| **Within** | | | |
| subject 1 | **p = 1.9e-04, q = 3.2e-04, t(59) = 4,<br>Cohen's dz = 0.45, SD: 0.085** | **p = 0.034, q = 0.046, t(59) = 2.2,<br>Cohen's dz = 0.24, SD: 0.069** | **p = 1e-07, q = 2.6e-07, t(59) = 6.1,<br>Cohen's dz = 0.68, SD: 0.08** |
| subject 2 | **p = 7.7e-09, q = 2.1e-08, t(59) = -6.7,<br>Cohen's dz = -0.75, SD: 0.13** | **p = 1.1e-12, q = 4.2e-12, t(59) = 9,<br>Cohen's dz = 1, SD: 0.11** | p = 0.24, q = 0.28, t(59) = 1.2,<br>Cohen's dz = 0.13, SD: 0.085 |
| subject 3 | **p = 7.2e-13, q = 3e-12, t(59) = -9.1,<br>Cohen's dz = -1, SD: 0.074** | **p = 2.8e-27, q = 2.9e-26, t(59) = 19,<br>Cohen's dz = 2.2, SD: 0.075** | **p = 1.1e-18, q = 6e-18, t(59) = 13,<br>Cohen's dz = 1.4, SD: 0.06** |
| subject 4 | **p = 4.1e-05, q = 7.3e-05, t(59) = -4.4,<br>Cohen's dz = -0.5, SD: 0.11** | **p = 1.1e-12, q = 4.3e-12, t(59) = 9,<br>Cohen's dz = 1, SD: 0.1** | **p = 3e-07, q = 7e-07, t(59) = 5.8,<br>Cohen's dz = 0.65, SD: 0.078** |
| **Mixed-Within** | | | |
| subject 5 | **p = 3.7e-26, q = 2.7e-25, t(59) = -18,<br>Cohen's dz = -2.1, SD: 0.1** | **p = 1.3e-26, q = 1.9e-25, t(59) = 19,<br>Cohen's dz = 2.1, SD: 0.1** | p = 0.22, q = 0.26, t(59) = -1.3,<br>Cohen's dz = -0.14, SD: 0.045 |
| subject 6 | **p = 7.5e-37, q = 1.4e-35, t(59) = -29,<br>Cohen's dz = -3.3, SD: 0.069** | **p = 1.2e-38, q = 2.9e-37, t(59) = 32,<br>Cohen's dz = 3.5, SD: 0.061** | **p = 0.024, q = 0.035, t(59) = -2.3,<br>Cohen's dz = -0.26, SD: 0.04** |
| **Mixed-Across** | | | |
| subject 7 | p = 0.43, q = 0.44, t(59) = -0.79,<br>Cohen's dz = -0.089, SD: 0.052 | p = 0.4, q = 0.42, t(59) = -0.85,<br>Cohen's dz = -0.095, SD: 0.042 | p = 0.066, q = 0.085, t(59) = -1.9,<br>Cohen's dz = -0.21, SD: 0.041 |
| subject 8 | p = 0.45, q = 0.45, t(59) = 0.77,<br>Cohen's dz = 0.086, SD: 0.043 | p = 0.046, q = 0.06, t(59) = -2,<br>Cohen's dz = -0.23, SD: 0.043 | p = 0.24, q = 0.28, t(59) = -1.2,<br>Cohen's dz = -0.13, SD: 0.046 |
| subject 9 | p = 0.37, q = 0.39, t(59) = 0.91,<br>Cohen's dz = 0.1, SD: 0.042 | **p = 0.0077, q = 0.012, t(59) = -2.8,<br>Cohen's dz = -0.31, SD: 0.041** | p = 0.1, q = 0.13, t(59) = -1.6,<br>Cohen's dz = -0.18, SD: 0.046 |
| **Across** | | | |
| subject 10 | p = 0.094, q = 0.12, t(179) = 1.7,<br>Cohen's dz = 0.12, SD: 0.04 | **p = 0.00011, q = 2e-04, t(179) = 3.9,<br>Cohen's dz = 0.28, SD: 0.039** | **p = 4.6e-07, q = 9.6e-07, t(179) = 5.2,<br>Cohen's dz = 0.37, SD: 0.043** |
| subject 11 | **p = 0.026, q = 0.037, t(179) = 2.2,<br>Cohen's dz = 0.16, SD: 0.055** | **p = 1.7e-14, q = 7.6e-14, t(179) = -8.4,<br>Cohen's dz = -0.59, SD: 0.044** | **p = 1.5e-07, q = 3.5e-07, t(179) = -5.5,<br>Cohen's dz = -0.39, SD: 0.044** |
| subject 12 | **p = 6.5e-06, q = 1.3e-05, t(179) = 4.6,<br>Cohen's dz = 0.33, SD: 0.043** | **p = 1.5e-08, q = 4e-08, t(179) = -5.9,<br>Cohen's dz = -0.42, SD: 0.04** | p = 0.26, q = 0.3, t(179) = -1.1,<br>Cohen's dz = -0.08, SD: 0.034 |

| S1 | Pre&Cue vs. Post&Cue | Post&Cue vs. Pre&Post | Pre&Cue vs. Pre&Post |
|---|---|---|---|
| **Within** | | | |
| subject 1 | p = 0.29, q = 0.32, t(59) = 1.1,<br>Cohen's dz = 0.12, SD: 0.1 | **p = 4.9e-11, q = 1.7e-10, t(59) = 8,<br>Cohen's dz = 0.9, SD: 0.069** | **p = 3.6e-07, q = 8.1e-07, t(59) = 5.7,<br>Cohen's dz = 0.64, SD: 0.12** |
| subject 2 | **p = 2.6e-10, q = 8.5e-10, t(59) = -7.6,<br>Cohen's dz = -0.85, SD: 0.075** | **p = 1.2e-16, q = 5.9e-16, t(59) = 11,<br>Cohen's dz = 1.3, SD: 0.073** | **p = 0.0017, q = 0.0027, t(59) = 3.3,<br>Cohen's dz = 0.37, SD: 0.08** |
| subject 3 | **p = 8.5e-10, q = 2.7e-09, t(59) = -7.3,<br>Cohen's dz = -0.82, SD: 0.099** | **p = 3.7e-26, q = 2.7e-25, t(59) = 18,<br>Cohen's dz = 2.1, SD: 0.09** | **p = 6.5e-20, q = 3.9e-19, t(59) = 14,<br>Cohen's dz = 1.5, SD: 0.068** |
| subject 4 | **p = 1.9e-09, q = 5.3e-09, t(59) = -7.1,<br>Cohen's dz = -0.79, SD: 0.079** | **p = 2.1e-20, q = 1.4e-19, t(59) = 14,<br>Cohen's dz = 1.6, SD: 0.077** | **p = 3.9e-12, q = 1.4e-11, t(59) = 8.7,<br>Cohen's dz = 0.97, SD: 0.06** |
| **Mixed-Within** | | | |
| subject 5 | **p = 9.1e-28, q = 1.1e-26, t(59) = -20,<br>Cohen's dz = -2.2, SD: 0.095** | **p = 2.1e-28, q = 3e-27, t(59) = 20,<br>Cohen's dz = 2.3, SD: 0.089** | p = 0.15, q = 0.18, t(59) = -1.5,<br>Cohen's dz = -0.16, SD: 0.035 |
| subject 6 | **p = 2.2e-44, q = 1.6e-42, t(59) = -40,<br>Cohen's dz = -4.5, SD: 0.057** | **p = 2.4e-43, q = 8.6e-42, t(59) = 38,<br>Cohen's dz = 4.3, SD: 0.055** | **p = 3.7e-07, q = 8.1e-07, t(59) = -5.7,<br>Cohen's dz = -0.64, SD: 0.026** |
| **Mixed-Across** | | | |
| subject 7 | p = 0.7, q = 0.7, t(59) = 0.39,<br>Cohen's dz = 0.044, SD: 0.033 | **p = 0.013, q = 0.02, t(59) = 2.6,<br>Cohen's dz = 0.29, SD: 0.036** | **p = 0.01, q = 0.015, t(59) = 2.7,<br>Cohen's dz = 0.3, SD: 0.039** |
| subject 8 | p = 0.32, q = 0.35, t(59) = -1,<br>Cohen's dz = -0.11, SD: 0.043 | p = 0.26, q = 0.3, t(59) = -1.1,<br>Cohen's dz = -0.13, SD: 0.045 | p = 0.043, q = 0.057, t(59) = -2.1,<br>Cohen's dz = -0.23, SD: 0.045 |
| subject 9 | **p = 0.016, q = 0.024, t(59) = 2.5,<br>Cohen's dz = 0.28, SD: 0.04** | **p = 2.1e-08, q = 5.5e-08, t(59) = -6.5,<br>Cohen's dz = -0.72, SD: 0.037** | **p = 4e-04, q = 6.7e-04, t(59) = -3.8,<br>Cohen's dz = -0.42, SD: 0.038** |
| **Across** | | | |
| subject 10 | **p = 1.2e-05, q = 2.2e-05, t(179) = 4.5,<br>Cohen's dz = 0.32, SD: 0.038** | **p = 1.7e-05, q = 3.9e-05, t(179) = 4.4,<br>Cohen's dz = 0.31, SD: 0.034** | **p = 4.3e-17, q = 2.2e-16, t(179) = 9.3,<br>Cohen's dz = 0.66, SD: 0.034** |
| subject 11 | **p = 1e-09, q = 3.1e-09, t(179) = 6.4,<br>Cohen's dz = 0.46, SD: 0.035** | **p = 7.3e-06, q = 1.4e-05, t(179) = -4.6,<br>Cohen's dz = -0.33, SD: 0.033** | **p = 0.033, q = 0.045, t(179) = 2.2,<br>Cohen's dz = 0.15, SD: 0.034** |
| subject 12 | **p = 0.008, q = 0.013, t(179) = 2.7,<br>Cohen's dz = 0.19, SD: 0.032** | p = 0.34, q = 0.37, t(179) = 0.95,<br>Cohen's dz = 0.067, SD: 0.031 | **p = 9.9e-07, q = 2e-06, t(179) = 5.1,<br>Cohen's dz = 0.36, SD: 0.023** |

### Supplementary 6: Single-subject statistical tests for corrected mean representational similarity of within-image comparisons

| V1 | Pre&Cue vs. Post&Cue | Post&Cue vs. Pre&Post | Pre&Cue vs. Pre&Post |
|---|---|---|---|
| **Within** | | | |
| subject 1 | *p* = 0.85, *q* = 1, t(19) = -0.2, Cohen's *dz* = -0.044, SD: 0.061 | **p = 1.9e-04, q = 0.002, t(19) = -4.6, Cohen's dz = -1, SD: 0.06** | **p = 7.9e-05, q = 0.0014, t(19) = -5, Cohen's dz = -1.1, SD: 0.058** |
| subject 2 | *p* = 0.12, *q* = 0.4, t(19) = 1.6, Cohen's *dz* = 0.37, SD: 0.088 | *p* = 0.36, *q* = 0.78, t(19) = -0.94, Cohen's *dz* = -0.21, SD: 0.077 | *p* = 0.2, *q* = 0.55, t(19) = 1.3, Cohen's *dz* = 0.3, SD: 0.054 |
| subject 3 | *p* = 0.9, *q* = 1, t(19) = 0.13, Cohen's *dz* = 0.028, SD: 0.041 | **p = 0.0052, q = 0.034, t(19) = -3.2, Cohen's dz = -0.71, SD: 0.058** | **p = 0.0081, q = 0.045, t(19) = -3, Cohen's dz = -0.66, SD: 0.06** |
| subject 4 | *p* = 0.72, *q* = 1, t(19) = -0.36, Cohen's *dz* = -0.081, SD: 0.053 | **p = 1.3e-04, q = 0.0019, t(19) = -4.8, Cohen's dz = -1.1, SD: 0.066** | **p = 1.4e-05, q = 5.1e-04, t(19) = -5.8, Cohen's dz = -1.3, SD: 0.058** |
| **Mixed-Within** | | | |
| subject 5 | *p* = 0.36, *q* = 0.78, t(19) = -0.94, Cohen's *dz* = -0.21, SD: 0.075 | *p* = 0.013, *q* = 0.066, t(19) = -2.7, Cohen's *dz* = -0.61, SD: 0.073 | **p = 4.5e-08, q = 3.2e-06, t(19) = -8.7, Cohen's dz = -2, SD: 0.031** |
| subject 6 | **p = 1.9e-04, q = 0.002, t(19) = -4.6, Cohen's dz = -1, SD: 0.039** | *p* = 0.85, *q* = 1, t(19) = 0.2, Cohen's *dz* = 0.044, SD: 0.059 | **p = 7.5e-05, q = 0.0014, t(19) = -5, Cohen's dz = -1.1, SD: 0.034** |
| **Mixed-Across** | | | |
| subject 7 | *p* = 0.51, *q* = 0.94, t(19) = -0.67, Cohen's *dz* = -0.15, SD: 0.035 | *p* = 0.046, *q* = 0.18, t(19) = -2.1, Cohen's *dz* = -0.48, SD: 0.04 | **p = 0.0058, q = 0.035, t(19) = -3.1, Cohen's dz = -0.69, SD: 0.035** |
| subject 8 | *p* = 0.28, *q* = 0.66, t(19) = -1.1, Cohen's *dz* = -0.25, SD: 0.047 | *p* = 0.083, *q* = 0.32, t(19) = -1.8, Cohen's *dz* = -0.41, SD: 0.039 | **p = 0.0023, q = 0.016, t(19) = -3.5, Cohen's dz = -0.79, SD: 0.035** |
| subject 9 | *p* = 0.17, *q* = 0.51, t(19) = -1.4, Cohen's *dz* = -0.32, SD: 0.03 | **p = 0.0022, q = 0.016, t(19) = -3.5, Cohen's dz = -0.79, SD: 0.034** | **p = 4.5e-04, q = 0.0041, t(19) = -4.2, Cohen's dz = -0.95, SD: 0.038** |
| **Across** | | | |
| subject 10 | *p* = 0.54, *q* = 0.94, t(19) = -0.63, Cohen's *dz* = -0.14, SD: 0.037 | *p* = 0.26, *q* = 0.65, t(19) = -1.2, Cohen's *dz* = -0.26, SD: 0.036 | *p* = 0.16, *q* = 0.51, t(19) = -1.5, Cohen's *dz* = -0.32, SD: 0.045 |
| subject 11 | *p* = 0.24, *q* = 0.64, t(19) = -1.2, Cohen's *dz* = -0.27, SD: 0.042 | *p* = 0.25, *q* = 0.64, t(19) = -1.2, Cohen's *dz* = -0.27, SD: 0.042 | *p* = 0.031, *q* = 0.13, t(19) = -2.3, Cohen's *dz* = -0.52, SD: 0.043 |
| subject 12 | *p* = 0.38, *q* = 0.78, t(19) = -0.9, Cohen's *dz* = -0.2, SD: 0.041 | *p* = 0.1, *q* = 0.37, t(19) = -1.7, Cohen's *dz* = -0.38, SD: 0.03 | *p* = 0.018, *q* = 0.087, t(19) = -2.6, Cohen's *dz* = -0.58, SD: 0.035 |

| S1 | Pre&Cue vs. Post&Cue | Post&Cue vs. Pre&Post | Pre&Cue vs. Pre&Post |
|---|---|---|---|
| **Within** | | | |
| subject 1 | *p* = 0.49, *q* = 0.93, t(19) = -0.7, Cohen's *dz* = -0.16, SD: 0.052 | *p* = 0.49, *q* = 0.93, t(19) = -0.71, Cohen's *dz* = -0.16, SD: 0.08 | *p* = 0.3, *q* = 0.7, t(19) = -1.1, Cohen's *dz* = -0.24, SD: 0.088 |
| subject 2 | *p* = 0.96, *q* = 1, t(19) = -0.047, Cohen's *dz* = -0.011, SD: 0.056 | *p* = 0.74, *q* = 1, t(19) = 0.34, Cohen's *dz* = 0.076, SD: 0.052 | *p* = 0.78, *q* = 1, t(19) = 0.29, Cohen's *dz* = 0.065, SD: 0.052 |
| subject 3 | *p* = 0.87, *q* = 1, t(19) = -0.17, Cohen's *dz* = -0.038, SD: 0.069 | *p* = 0.78, *q* = 1, t(19) = -0.28, Cohen's *dz* = -0.063, SD: 0.063 | *p* = 0.58, *q* = 0.98, t(19) = -0.56, Cohen's *dz* = -0.12, SD: 0.053 |
| subject 4 | *p* = 0.43, *q* = 0.87, t(19) = -0.8, Cohen's *dz* = -0.18, SD: 0.046 | *p* = 0.38, *q* = 0.78, t(19) = 0.9, Cohen's *dz* = 0.2, SD: 0.047 | *p* = 0.89, *q* = 1, t(19) = 0.13, Cohen's *dz* = 0.03, SD: 0.041 |
| **Mixed-Within** | | | |
| subject 5 | *p* = 0.93, *q* = 1, t(19) = 0.093, Cohen's *dz* = 0.021, SD: 0.091 | *p* = 0.92, *q* = 1, t(19) = 0.1, Cohen's *dz* = 0.022, SD: 0.091 | *p* = 0.55, *q* = 0.94, t(19) = 0.61, Cohen's *dz* = 0.14, SD: 0.029 |
| subject 6 | *p* = 0.86, *q* = 1, t(19) = 0.18, Cohen's *dz* = 0.041, SD: 0.052 | *p* = 0.98, *q* = 1, t(19) = -0.025, Cohen's *dz* = -0.0056, SD: 0.062 | *p* = 0.75, *q* = 1, t(19) = 0.33, Cohen's *dz* = 0.073, SD: 0.024 |
| **Mixed-Across** | | | |
| subject 7 | *p* = 1, *q* = 1, t(19) = -0.0034, Cohen's *dz* = -7.5e-04, SD: 0.024 | *p* = 0.83, *q* = 1, t(19) = -0.22, Cohen's *dz* = -0.049, SD: 0.033 | *p* = 0.8, *q* = 1, t(19) = -0.26, Cohen's *dz* = -0.058, SD: 0.028 |
| subject 8 | *p* = 0.53, *q* = 0.94, t(19) = 0.63, Cohen's *dz* = 0.14, SD: 0.033 | *p* = 0.2, *q* = 0.55, t(19) = -1.3, Cohen's *dz* = -0.3, SD: 0.024 | *p* = 0.73, *q* = 1, t(19) = -0.35, Cohen's *dz* = -0.079, SD: 0.033 |
| subject 9 | *p* = 0.67, *q* = 1, t(19) = -0.43, Cohen's *dz* = -0.096, SD: 0.037 | *p* = 0.99, *q* = 1, t(19) = -0.017, Cohen's *dz* = -0.0037, SD: 0.035 | *p* = 0.69, *q* = 1, t(19) = -0.41, Cohen's *dz* = -0.092, SD: 0.04 |
| **Across** | | | |
| subject 10 | *p* = 0.024, *q* = 0.11, t(19) = 2.5, Cohen's *dz* = 0.55, SD: 0.022 | *p* = 0.14, *q* = 0.46, t(19) = -1.5, Cohen's *dz* = -0.34, SD: 0.029 | *p* = 0.73, *q* = 1, t(19) = 0.35, Cohen's *dz* = 0.079, SD: 0.025 |
| subject 11 | *p* = 0.96, *q* = 1, t(19) = 0.055, Cohen's *dz* = 0.012, SD: 0.035 | *p* = 0.73, *q* = 1, t(19) = 0.35, Cohen's *dz* = 0.077, SD: 0.036 | *p* = 0.74, *q* = 1, t(19) = 0.34, Cohen's *dz* = 0.076, SD: 0.04 |
| subject 12 | *p* = 0.77, *q* = 1, t(19) = -0.29, Cohen's *dz* = -0.065, SD: 0.032 | *p* = 0.91, *q* = 1, t(19) = 0.11, Cohen's *dz* = 0.024, SD: 0.033 | *p* = 0.82, *q* = 1, t(19) = -0.23, Cohen's *dz* = -0.05, SD: 0.025 |
